# Reconstructing plant beneficial bacterial consortia by integrating dilution-to-extinction microbiome perturbation with genome-resolved synthetic ecology

**DOI:** 10.64898/2026.04.09.717421

**Authors:** Jiayi Jing, Adam Ossowicki, Vittorio Tracanna, Elio G. W. M. Schijlen, Mirna L. Baak, Walter Pirovano, Wilfred F.J van Ijcken, Dominika Rybka, Saskia Gerards, Somayah S. Elsayed, Zachary L Reitz, Gilles P. van Wezel, Jos M. Raaijmakers, Paolina Garbeva, Marnix H. Medema

**Author notes:** Correspondence to: Marnix H. Medema, Bioinformatics Group, Wageningen University, Droevendaalsesteeg 1, 6708PB, Wageningen. Mail; Paolina Garbeva, Netherlands Institute of Ecology (NIOO-KNAW), Department of Microbial Ecology, Droevendaalsesteeg 10, 6708 PB, Wageningen, The Netherlands. Mail. These authors contributed equally to this work (Adam Ossowicki, Vittorio Tracanna).

## Abstract

Across the biosphere, microbiomes play essential roles in shaping the health of their host. One notable example of such a microbiome-associated phenotype is disease-suppressive soils, where susceptible plant hosts enrich and activate specific rhizosphere microbial consortia for protection against fungal root pathogens. However, identifying and reconstructing microbial consortia responsible for host protection remains challenging, given the inherent taxonomic and functional complexity of microbiomes. Here, we integrated metagenomic profiling of disease-suppressive microbiomes perturbed by dilution-to-extinction (DTE) with comprehensive culturing and synthetic ecology to identify the key bacterial taxa conferring suppressiveness to the fungal wheat pathogen *Fusarium culmorum*. Metagenomics of wheat rhizosphere samples along the DTE trajectory revealed bacterial taxa and functions associated with the disease-suppressive phenotype. Crosslinking these DTE metagenome data with a genome-sequenced collection of 336 rhizobacterial isolates from the suppressive soil allowed the reconstruction of synthetic communities (SynComs) of 11 de-replicated strains negatively associated with disease severity. Upon re-introduction in sterilized suppressive soils, this SynCom consistently reproduced the disease-suppressive phenotype. Paired time-series metagenomics and metatranscriptomics of the SynComs pinpointed candidate biosynthetic gene clusters, including a novel non-alpha poly-amino-acid (NAPAA) gene cluster from *Arthrobacter*, upregulated in presence of *F. culmorum*. Chemically synthesized NAPAA variants ε-poly-_L_-lysine and δ-poly-_L_-ornithine significantly inhibited *F. culmorum* hyphal growth. Collectively, our work establishes a transformative strategy for reconstructing microbial consortia that recapitulates beneficial microbiome-associated phenotypes in plant and animal kingdoms.

## Introduction

Microbiomes shape host health across all domains of life^1,2^, yet identifying the specific microbial taxa and functional pathways responsible for these phenotypes remains challenging owing to the complex, multi-species nature of ecosystems, which can obscure the contributions of individual members. Soil ecosystems exemplify this challenge through their exceptional microbial diversity^3,4^, particularly in ‘disease-suppressive’ soils, which protect susceptible host plants against bacterial or fungal pathogens^5^. For most disease-suppressive soils, protection is mediated by the activities of specific members of the soil and rhizosphere microbiome^6–8^. Deciphering the mechanisms underlying disease-suppressiveness is instrumental for developing new approaches to combat soil-borne diseases which threaten global food security and food safety^9–11^. Recent studies across diverse crops have documented pathogen suppression through distinct microbiome-mediated mechanisms such as iron competition^12^, antibiotic production^13,14^, and induced systemic resistance^15,16^.

Efforts to simplify such complex systems have generally followed either top-down approaches through perturbations^17^ such as heat^18,19^, biocides^20^, and dilutions^21,22^, or bottom-up approaches that reconstruct minimal communities from cultured microbial taxa with predicted/tested functions or abundance patterns^23–25^. However, both approaches have inherent limitations. Top-down approaches typically rely on 16S/functional amplicon sequencing and/or shotgun metagenomics, which remain descriptive and are unable to resolve individual functional contributions to complex phenotypes. Conversely, bottom-up synthetic approaches, while offering precise compositional control, may not reflect mechanisms active in the native soil microbial communities. These limitations have precluded the comprehensive identification and validation of the strains, genes, and mechanisms underlying complex microbiome-mediated phenotypes such as disease suppression.

Here, we combine and integrate bottom-up and top-down approaches to identify taxa and genes that are likely causally associated with disease suppression in wheat rhizosphere microbiomes. Through large-scale screening of 28 agricultural field soils, we previously identified a soil (S11) with robust microbiome-mediated suppressiveness against the fungal pathogen *Fusarium culmorum* in wheat^26^. Disease-suppressiveness of soil S11 to *F. culmorum* could be eliminated by selective heat treatments and transplanted by mixing 10% of soil S11 into conducive soils^26^. While comparative profiling of suppressive and conducive soils revealed few consistent taxonomic signatures, functional amplicon sequencing of adenylation domains (A domains) revealed that soil S11 harbours diverse biosynthetic gene clusters (BGCs) of interest^27^. To identify the active members within soil S11, we combined top-down dilution-to-extinction (DTE) with bottom-up synthetic community (SynCom) design, underpinned by a genome sequenced collection of 336 rhizosphere bacterial isolates from S11 soil. By correlating the loss of suppressiveness along the DTE trajectory with isolate-resolved (meta)genomic profiles, we identified 11 bacterial strains that, when reconstructed as a SynCom, fully recapitulate disease-suppressive phenotype. Crucially, by using the bacterial strains’ genomes as high-resolution references for metatranscriptomic analysis, we identified a novel, pathogen-induced non-alpha poly-amino-acid (NAPAA) BGC carried by a strain present in relatively low abundance in the SynCom (*Arthrobacter equi*). We further demonstrate that NAPAA variants, including ε-poly-_L_-lysine and δ-poly-_L_-ornithine, exhibit potent antifungal activity against *F. culmorum in vitro*. Our results show that disease suppressiveness of soil emerges from coordinated regulation across phylogenetically distinct taxa, highlighting the necessity of strain-resolved experimental and computational frameworks to decode the rare but functionally pivotal members of the beneficial microbiome.

## Results

### Dilution-to-extinction progressively dilutes microbial community complexity and diminishes disease-suppressiveness

To systematically identify bacterial taxa associated with soil S11’s disease suppressiveness, we employed dilution-to-extinction (DTE) to progressively reduce microbiome complexity and reveal essential microbial contributors through metagenomic analysis. We extracted the rhizosphere microbiome of wheat plants grown for two weeks in soil S11 and introduced serial dilutions of the rhizosphere microbiome (10X - 400X) into sterile sandy soil. The extracted soil microbiome (ESM) conferred significant protection against *F. culmorum* root infection, with suppressive activity decreasing non-linearly: low dilutions (10X - 50X) maintained suppression, while higher dilutions (100X - 400X) lost this suppressive effect (Fig.1a-c).

**Fig. 1.**
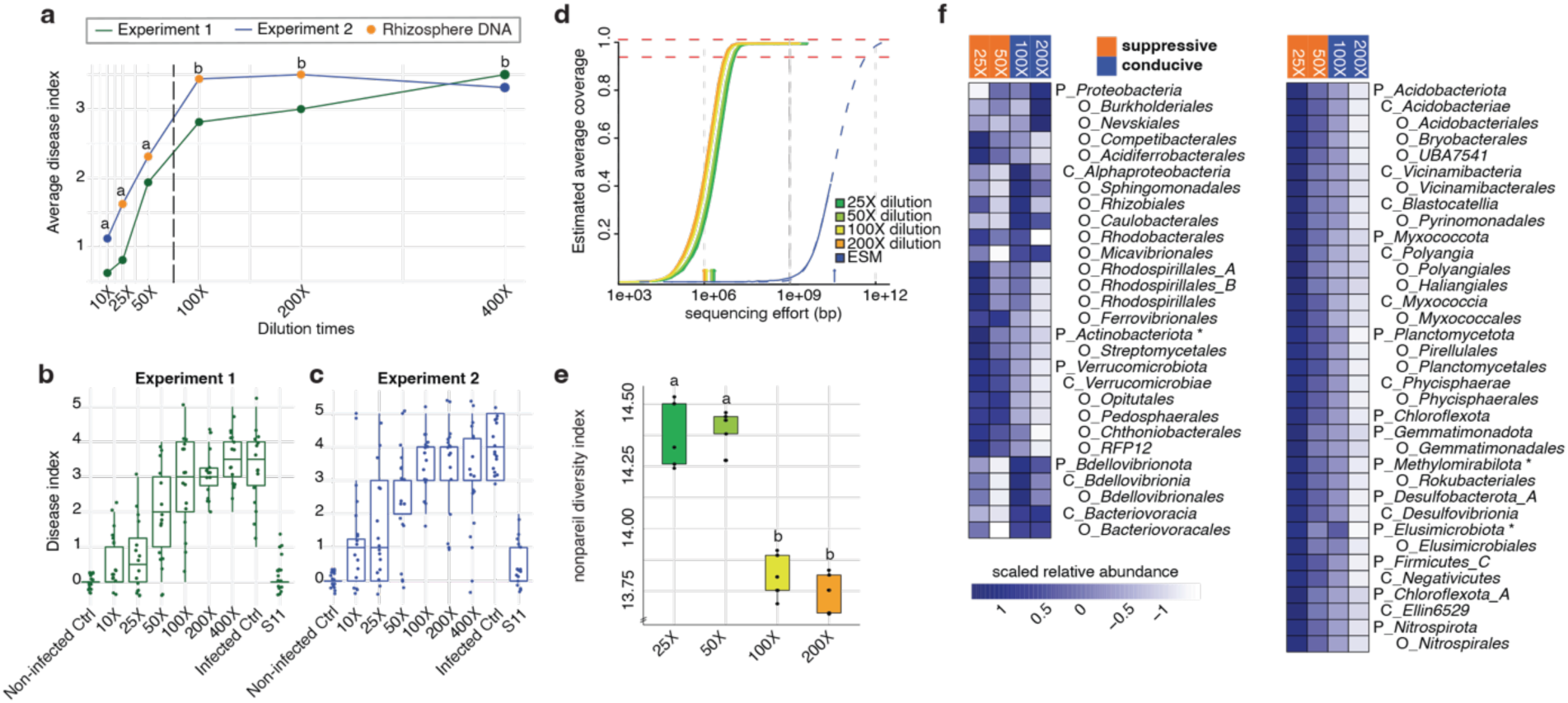
Dilution-to-extinction dissects microbial determinants of disease suppression in the wheat rhizosphere. (**a**) Combined results from two independent dilution-to-extinction experiments with *Fusarium*-suppressive soil S11. The black dash line indicates the transition point from disease-suppressive to conducive phenotypes as determined by chi-square statistical analysis. Letters above data points represent statistically significant differences among dilutions. Metagenomics analyses were conducted using rhizosphere DNA samples collected from four dilutions (orange dots) in experiment 2. **(b-c)** Boxplots illustrating disease severity from two independent experiments. Non-infected Ctrl: sterile soil without pathogen inoculation; Infected Ctrl: sterile soil with pathogen inoculation; S11: pathogen-induced suppressive field soil. **(d)** Estimated sequencing coverage (ESM) of microbial diversity in the original suppressive soil S11 extract and rhizosphere samples derived from different ESM dilutions. **(e)** Impact of introduced S11 into sterile soil on genetic diversity of wheat rhizosphere microbiomes, measured by Nonpareil diversity index. Different letters indicate significant differences (ANOVA and Tukey’s post-hoc test, p < 0.001) among groups. **(f)** Significant shifts in relative abundance of bacterial taxa between rhizosphere microbiomes from *Fusarium*-suppressive (25X and 50X) versus non-suppressive (100X and 200X) dilutions of S11. Abundance is scaled by rows (z-score). Taxonomic levels are indicated by letters: P, phylum; C, class; O, order. Statistical significance assessed by DESeq2 (adjusted p-value < 0.05). Taxa marked with an asterisk are **<u>not</u>** statistically significant but indicate the phylum of subordinate taxa. Low-abundance taxa (<1% for all samples) were omitted for clarity.

We performed shotgun metagenomic sequencing of rhizosphere samples from four ESM dilutions spanning suppressive to conducive phenotypes (orange dots in Fig.1a), achieving near-complete coverage of the microbiota (Fig.1d). While taxonomic profiles remained consistent across biological replicates (Fig.S1), microbial diversity decreased with increasing dilution (Fig.1e), with the Nonpareil diversity index shifting from approximately 14.5 to 13.8, correlating with the loss of disease suppression. Bacterial community composition showed dilution-dependent shifts: *Proteobacteria* and *Bacteroidota* dominated (together constituting 60% to 68% of the bacterial community) and increased at higher ESM dilutions (Fig.1f). While the relative abundance of these dominant taxa (including most underlying genera) remained stable or even increased, several lower-abundance taxa declined significantly, showing a concomitant reduction in the suppressive phenotype (Fig.1f). Metagenomic profiling based on assembled contigs and metagenome-assembled-genomes (MAGs) identified specific BGCs and functional orthologs, such as Type VI secretion system (T6SS), iron uptake and chitinases, that were significantly enriched in suppressive soils (Supplementary Results, Fig.S2, Table S1-S2). While these top-down analyses identified a core set of suppression-associated functions, we next sought to capture this diversity through a comprehensive culture collection to recapitulate the suppressive phenotype *in planta*.

### Genome-resolved associations guide design and reconstitution of suppressive SynComs

To enable strain-level resolution, we established a comprehensive culture collection from 10X ESM by utilizing diverse media and growth conditions, obtaining 336 initial isolates spanning 5 phyla and 19 orders (Table.S3). We then performed multiplexed Illumina whole-genome sequencing of the entire strain collection and bioinformatic dereplication based on 40 universal marker genes^28^ to consolidate these isolates into 244 high-quality, unique taxonomic strain-representative draft genomes (with a median N50 of 21.5 kb, Table. S4). The strain collection represented approximately 40 % - 60% (with a median of 49.42%) of the bacterial reads from DTE samples and 45 % - 70 % (median of 57.39%) of the DTE metagenome assembled contigs (Table. S5). Comparative analyses revealed that isolate genomes encompassed a substantial portion of microbial diversity at finer taxonomic scales that were not captured by DTE-MAGs (Fig.S3). The metagenomics reads from the DTE experiment were mapped to these genomes and through a correlation coefficient (CC) analysis using abundance and disease indices of 20 DTE samples, we identified 11 bacterial strains (after de-replication) that showed significant negative correlation with disease severity (linear regression, p<0.05, N = 20; Fig.S4). They were taxonomically delineated as *Arthrobacter equi*, *Pedobacter* sp., *Mucilaginibacter* sp., *Pseudomonas* sp., *Amnibacterium* sp., *Plantibacter cousiniae*, and *Microbacterium* sp. (Fig.2a-b), most of which have previously been associated with wheat health and/or disease suppression^29–33^.

**Fig. 2.**
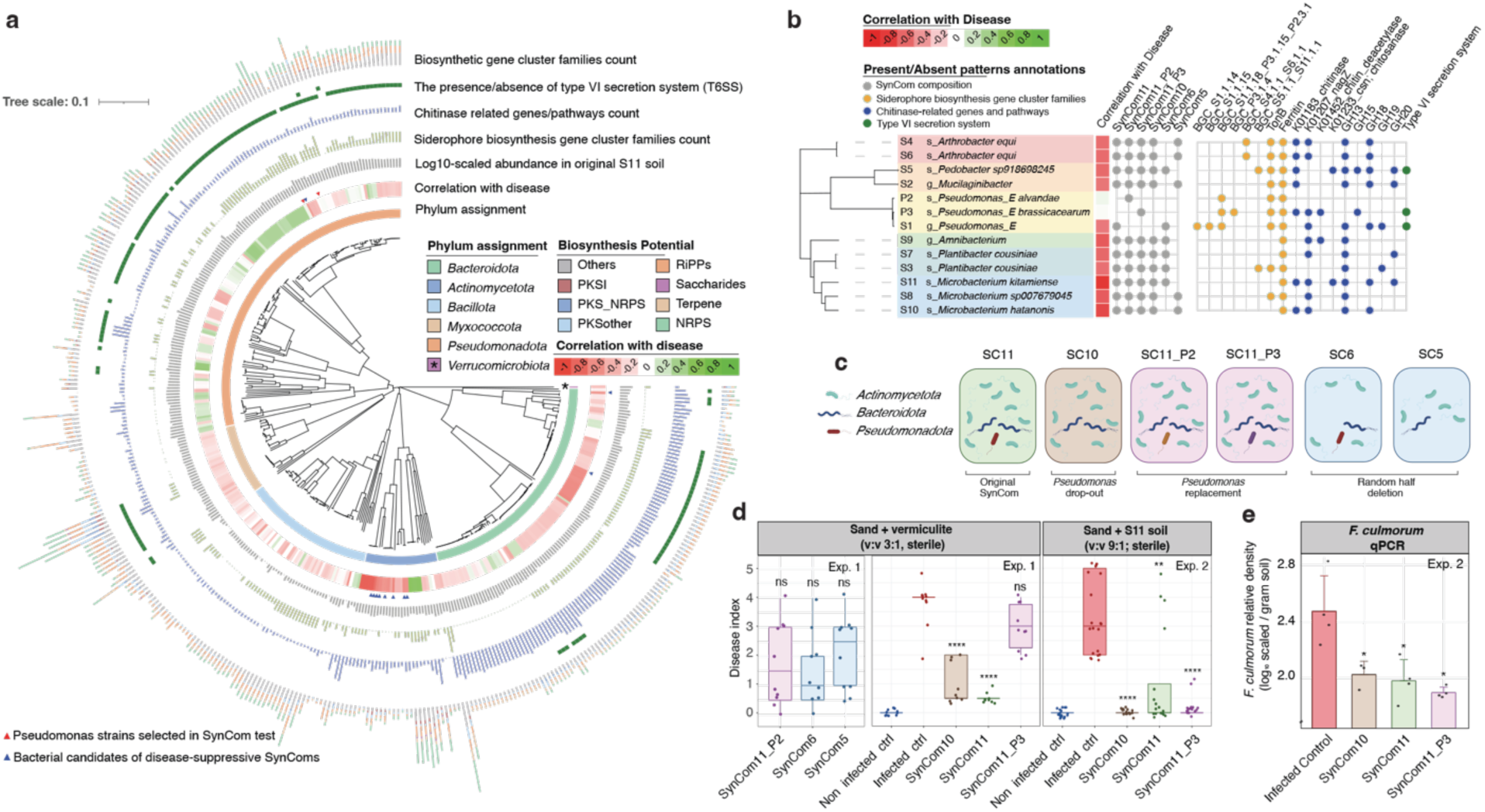
Correlation-guided synthetic communities design for functional validation. (**a**) Bootstrapped ML tree of 356 bacterial isolates cultured from the 10X dilution rhizosphere sample in the dilution-to-extinction experiment. From the inner to outer circles, the tree displays: phylum assignment, correlation with disease index, log10-scaled abundance in the original suppressive soil S11, number of siderophore biosynthetic gene clusters (BGCs), count of chitinase-related genes/pathways, presence/absence of type VI secretion system (T6SS), and number of biosynthetic gene cluster families (GCFs). BGC types were color-coded. Next, isolates were associated with a color gradient from red (positively correlated, conducive-associated) to green (negatively correlated, suppressive-associated) indicated the correlation with disease severity. Blue triangles indicate suppressive-associated bacterial isolates which were selected for synthetic communities (SynComs) construction. The red triangles highlight two additional *Pseudomonas* strains that were included in the replacement SynComs test. **(b)** Functional annotation of the 13 selected SynCom candidates, showing presence of siderophore BGCs (orange), chitinase-related genes/pathways (blue), and T6SS components (green). **(c)** Design strategy of the synthetic communities. SynCom11 comprised all (de-replicated) suppressive-associated isolates. A drop-out version (SynCom10) excluded the *Pseudomonas* strain from SynCom11. Two replacement SynComs (SynCom11_P2, SynCom11_P3) substituted the *Pseudomonas* S1 isolate with strains P2 and P3, respectively. Two randomly reduced communities (SynCom6 and SynCom5) included half of the SynCom11 members for evaluating community stability and performance. **(d)** Plant phenotype outcomes from two independent SynCom experiments. Three SynComs (SynCom11_P2, SynCom6, SynCom5) were tested in only experiment 1 (sand/vermiculite, 3:1, v/v, sterile) and observed big variation within the same group. Three SynComs (SynCom10, SynCom11, SynCom11_P3) were tested two different soil condition (sandy soil/S11, 9:1 v/v, sterile) to check their repeatability and collect rhizosphere DNA, RNA for sequencing. Statistical significance compared to the infected control were assessed by ANOVA with Tukey’s test: p<0.01 (“**”), and p<0.0001(“****”). Non-infected control plants were grown without *F. culmorum*; infected control plants and SynCom-treated plants were inoculated with *F. culmorum*. **(e)** The bar plots shown the quantification of *F. culmorum* in the rhizosphere soil samples. Fungal density was measured by qPCR, normalized by rhizosphere soil weight, and plotted on a log10-scale. Statistical analyses were performed for each treatment compared to infected control and were assessed by ANOVA with Tukey’s test: p<0.05 (“*”), and p<0.01 (“**”).

Genomic annotation of the strain collection revealed diverse functional genes and gene clusters (Fig.2a). The log_10_-scaled abundance of these bacterial strains in the S11 metagenome obtained in our previous study^26^, showed these strains were all highly abundant (log_10_-scaled RPM 3.06 - 4.68) in the original S11 disease-suppressive soil rhizosphere metagenome (Table. S6). According to the functional assignment, most of these prioritized 11 bacterial strains harbour at least 2 of the 3 functions pinpointed in our DTE-MAG analyses (Supplementary Results, Fig.S2), i.e. siderophore-, chitinase-, or T6SS- related functions. They were, however, not the strains with the largest numbers of BGCs, suggesting that the correlation between BGCs counts and their putative disease-suppressive capability is not linear. For example, in the *Actinomycetota* clade, the bacterial strains with higher BGC counts even showed a positive correlation with disease. Within the *Pseudomonadota* clade, there were only few bacterial strains that showed a negative correlation to disease and only *Pseudomonas* strain S1 is significant negatively correlated (Fig.2a). To further explore the functional differences within these taxonomically closely related *Pseudomonas* strains, we selected two (referred to as P2, and P3, indicated in Figure 2a with a red triangle) for a replacement assay to validate their disease-suppressive effects. Strain P2 is closest to strain S1, and P3 is the only strain containing the BGC for 2,4-diacetylphloroglucinol (2,4-DAPG) (Fig.S5), a well-known polyketide involved in disease suppressiveness to take-all disease of wheat^34^ and suppression of other fungal root diseases^35–37^. Additionally, we found that all the selected 11 strains exhibited intermediate predicted growth rates, with predicted doubling times between 1 and 3h based on codon usage bias analysis of the genomes (using gRodon R package)^38^. In contrast, both fast growers (predicted doubling time <1h) and slow growers (>3h) showed only weak correlations with disease severity (Pearson’s r from −0.5 to 0.5), suggesting that the DTE perturbation approach does not necessarily select for only fast-growing strains (Fig.S6).

In order to assess whether combinations of these 11 strains would be able to recapitulate the disease-suppressive phenotype, we designed several SynComs including: **i)** the 11 prioritized strains (SynCom11); **ii)** the drop-out SynCom (SynCom10) lacking *Pseudomonas* strain S1; **iii)** two replacement SynComs (SynCom11_P2, SynCom11_P3) where S1 is replaced with P2 and P3; and **iv)** two randomly depleted SynComs (SynCom6, SynCom5) (Fig.2c). The detailed compositional assignment of these SynComs and their taxonomic and functional annotations are shown in Fig.2b. In experiment 1, we tested the six SynComs and three individual *Pseudomonas* strains and observed that SynCom11 and SynCom10 showed robust suppression of *F. culmorum* of wheat compared to the control (Fig.2d, Fig.S7). SynCom11_P2, SynCom5, and SynCom6 revealed large variations across replicates and SynCom11_P3 did not provide any protection (Fig.2d). To check reproducibility and consistency of the disease-suppressive effects, we repeated the bioassay (experiment 2) with SynCom10, SynCom11, and SynCom11_P3 in a different soil condition (sandy soil: S11 soil, 9:1 (v/v), both sterile) optimized for high-quality RNA and DNA extractions for qPCR-based quantification of *F. culmorum* density and for metatranscriptomics. The three SynComs tested have only one strain difference but revealed different phenotypic outcomes (Fig.2d). From these two independent experiments, we found that SynCom10 provides the most robust and consistent disease suppressiveness across the two soil conditions, whereas SynCom11 and SynCom11_P3 revealed soil-type dependent effects and/or showed more variation across replicates. The qPCR analyses of *F. culmorum* further revealed significant reduction of pathogen density in the rhizosphere of all three SynCom treatments when 10% sterile S11 soil was introduced (Fig 2e). These findings confirm that the DTE-based correlation analysis is a robust approach for identifying and reconstructing disease-suppressive bacterial consortia.

### Pathogen-induced transcriptional shifts reveal trade-offs between community and strain-specific defense

We next performed integrative time-series metagenomic and metatranscriptomic analyses using high-quality genomes as references. Reference genomes were obtained through nanopore sequencing, assembling and further polishing using the short reads from Illumina sequencing, seven genomes of the SynCom members were assembled into single contigs, four into 2-5 contigs, and only the *Amnibacterium* (S9) genome remained genomically complete but fragmented (Table. S7). Rhizosphere profiling revealed that community structure remained stable between day 10 and day 20 under pathogen challenge, and notably, the introduction of *Pseudomonas* (S1 or P3) did not strongly affect the relative abundance of the other ten strains (Fig. 3b). While *Pedobacter* (S5) and *Mucilaginibacter* (S2) dominated in abundance, their transcriptomes exhibited comparatively low overall expression levels; conversely, the *Arthrobacter* strains (S4 and S6) were among the least abundant but most transcriptionally active, harboring the largest number of highly expressed genes (RNA TPM > 100) (Fig. S8d, Fig. S9). Metatranscriptomic analysis identified day 10 as a period of intensified microbial interaction, with SynCom10 exhibiting more than twice the number of pathogen-induced differentially expressed genes (DEGs) compared to *Pseudomonas*-containing consortia (Fig.3c). These DEGs were distributed across diverse taxa and included the widespread upregulation of ABC transporter genes, specifically the efflux-related *drrA* and *drrB* (in strains S3, S7, S9, S10), which likely mediate antibiotic resistance or cellular detoxification and may thus mediate defense against fungal secondary metabolites^39^, (Fig.S10, Fig.S11) alongside a GH18 chitinase in *Microbacterium kitamiense* strain S11 (Fig. S12c).

**Fig. 3.**
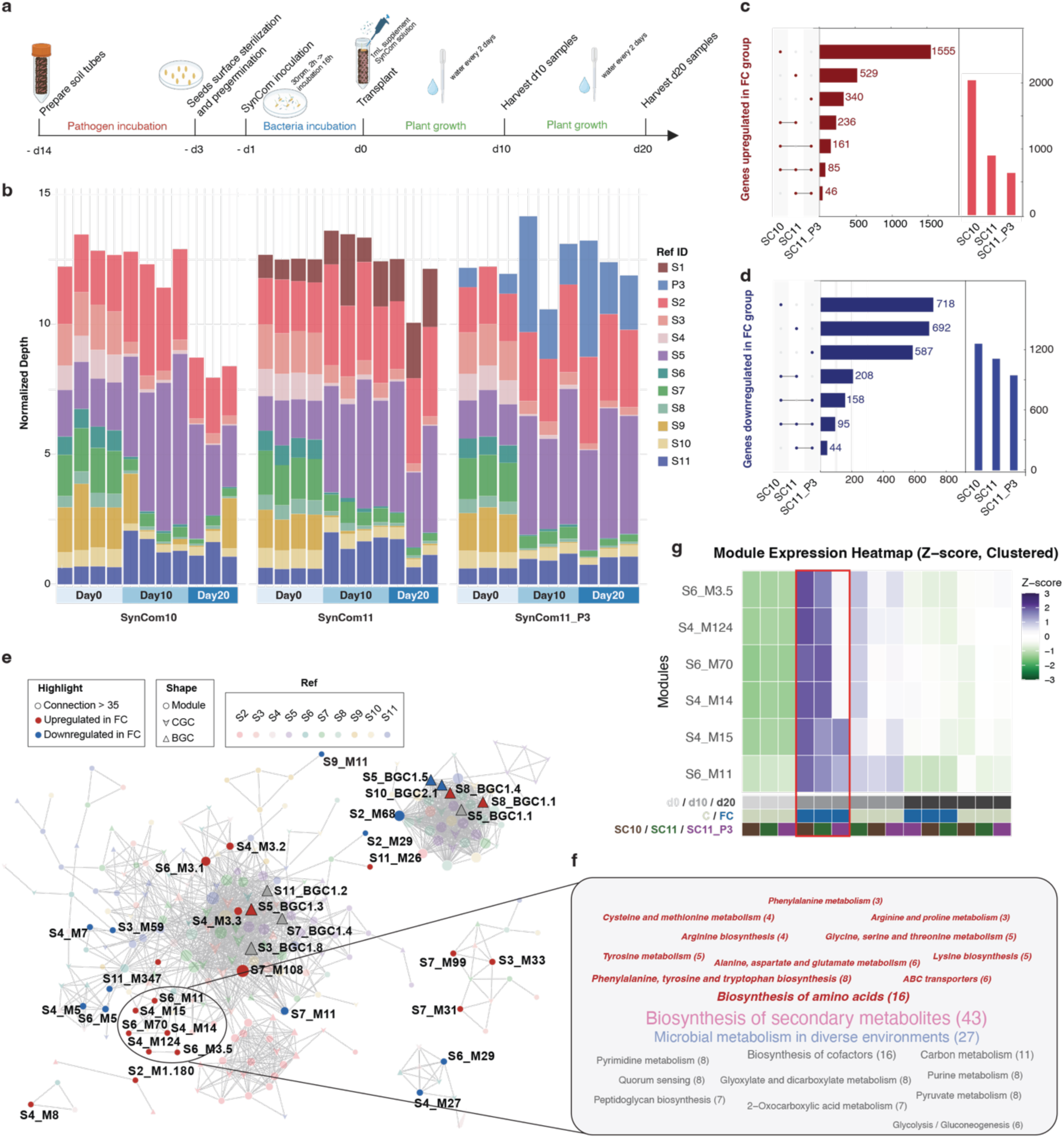
Compositional and transcriptional changes in SynComs in response to the fungal root pathogen. **(a)** The timeline of SynCom assay experiment 2. **(b)** The bar plots show the normalized depth of the inoculated bacteria in the rhizosphere of 3 SynComs under pathogen stress based on shotgun metagenomics data. Corresponding community composition data for non-infected controls were shown in Fig. S8. Color-bar from light to dark blue represents DNA samples collected at day 0, day 10, and day 20, respectively. Each color indicates a different bacterial member of SynCom. Day 0 samples represent the initial inoculum, while day 10 and day 20 samples were collected from rhizosphere soil. **(c - d)** Upset plots showing shared and unique up-regulated (d) and down-regulated (e) differentially expression genes (DEGs) across 3 SynComs conditions. DEGs were filtered by adjusted p-value <0.05 (DESeq2). Red bars represent up-regulated DEGs in *Fusarium*-inoculated groups versus non-*Fusarium*-inoculated groups; blue bars indicate the down-regulated DEGs. These plots were based on samples from day10 groups. **(e)** The global co-expression network (Pearson r > 0.9) shows co-regulated multigene modules, BGCs, and catabolic gene clusters (CGCs) across all samples. The analysis was streamlined by collapsing co-expressed modules (Table.S9) and using key functional terms (including BGCs and catabolic gene clusters [CGCs] in Table.S10&11) as individual network nodes. The round nodes represent modules which were defined based on co-expression pattern across different samples. The triangle nodes represent BGCs, and the V-shapes represent CGCs. Node size corresponds to the number of connected edges. The highlights show: red color - *Fusarium*-induced functions; blue color - *Fusarium*-suppressed functions; black boarder - BGCs with more than 40 connections. The rest non-highlighted node color was coding by their reference bacteria. **(f)** The KEGG pathways assignment of clustered *Fusarium*-induced modules. The pathway related to biosynthesis of secondary metabolites, transport, and metabolism of amino acid were highlighted in red color and bold font. **(g)** The heatmap of the expression of selected modules in different groups. The expression was based on median RNA TPM and scaled by per module with Z-score. The color bar below the heatmap indicated which different conditions including harvest date, pathogen stress, and SynComs treatment. The red board highlights the day 10 samples under pathogen stress from 3 SynCom-inoculated rhizosphere samples.

Microbial activity varied significantly over time, with day 20 comparisons demonstrating a significant suppression of pathogen transcriptional activity in all SynCom-treated plants relative to controls (Fig. S8e, f). The presence of *Pseudomonas* P3 shifted the suppressive landscape; in the absence of the pathogen, P3 dominated the community and the transcriptional activity of other members was suppressed (7 of 10 genomes show lower number of genes with RNA TPM > DNA TPM when P3 strain was present in the communities), a trend that was partially attenuated when the pathogen was present (Fig. S8a-c, Fig. S13, Table. S8). Detailed analysis of the *Pseudomonas* P3 genome revealed a dual antimicrobial strategy: the 2,4-DAPG BGC was constitutively active at high levels (TPM > 500), whereas the hydrogen cyanide (HCN) BGC was specifically induced by *F. culmorum* (Fig. S14). While ternary plots showed broadly similar transcriptional responses across all three SynComs, the subtle divergence in SynCom11_P3 suggests that P3’s dominance may impose a trade-off between community-wide cooperative and individual strain-driven defense (Fig. S15). These results pinpoint day 10 as the critical window of pathogen-microbiome interaction, highlighting a shift from broad community responses in SynCom10 to a more localized, specialized defense in the presence of *Pseudomonas* P3.

To bridge the top-down metagenomic predictions from DTE analysis with our bottom-up SynCom models, we validated the expression of suppression-associated features through a four-layer integrative analysis (Supplementary Results; Table S1-S2, Fig. S16a–e). While pathways such as Type IV pili, T6SS, and branched chain amino acid (BCAA) biosynthesis were significantly activated in the suppressive SynCom rhizosphere in the presence of *F. culmorum*, mechanisms like siderophore-mediated iron competition and chitinase activity lacked consistent pathogen-induced responses (Supplementary Results; Fig.S12, Fig.S16f, Fig.S17-18), suggesting that the core suppressive function may rely on previously uncharacterized molecular strategies.

### Genes associated with amino acid biosynthesis and nitrogen metabolism are enriched in disease-suppressive SynCom

To help elucidate the SynCom mechanisms underlying disease suppression and to identify functionally interacting gene modules across strains, we performed a global co-expression network analysis, (Fig.3e) revealing coordinated expression across community members. While some co-expression was observed among phylogenetically close species (i.e., S4/S6 and S3/S7), the predominant feature was the largest cluster spanning multiple diverse strains, highlighting a coordinated, SynCom transcriptional response to pathogen challenge. With this cluster, we observe highly connected BGCs (with more than 35 edges) and pathogen-induced functions that showed extensive co-expression across multiple categories and strains, suggesting they represent core responses. Among the highlighted modules, we identified a cohesive cluster of co-expression modules (S4_M14, S4_M15, S6_M11, S6_M70, S6_M3.5, and S4_M124) that are induced in the presence of the *F. culmorum*. Interestingly, many of the KOs from these modules are related to the biosynthesis, metabolism, and transport of amino acids (Fig.3f, Table.S12&13), reinforcing the previously identified importance of the BCAA biosynthesis pathway (Fig.S18). Another prominent annotation among these genes/transcripts is biosynthesis of secondary metabolites (Fig. 3f). As shown by the heatmap, these modules exhibited elevated expression levels in *Fusarium*-inoculated rhizosphere samples at day 10, with the strongest upregulation observed in SynCom10 and SynCom11 (Z-scores >1 for all 6 modules) compared to control conditions (Fig.3g). Metatranscriptomic profiling of the *Fusarium*-inoculated wheat rhizosphere further identified *Arthrobacter equi* and *Plantibacter cousiniae* as primary drivers of nitrogen metabolism and stress response (Supplementary Results; Fig.S19&Fig.S9), whereas *Pseudomonas* members were characterized by highly active biofilm formation functions in both SynCom11 and SynCom11_P3 (Supplementary Results; Fig.S20).

### A novel non-alpha poly-amino-acid (NAPAA) biosynthetic gene cluster (BGC) is associated with disease suppressiveness

Beyond the metabolic shifts, we next investigated the community’s capacity for producing complex specialized metabolites in response to the presence of the pathogen by analysing BGC expression patterns. The BGC annotations revealed a diverse set of clusters across 12 SynCom genomes (Table. S14), including those encoding the production of RiPPs (e.g., targeted antimicrobials), terpenes (e.g., membrane disruptors), nonribosomally synthesized peptides (NRPs, e.g., antifungal peptides), metallophores (facilitating iron competition with pathogen) and more. We assessed BGC expression at a global scale, focusing on the core biosynthesis genes within each BGC (full results in Fig.S21.; highlighted examples in Fig.4a). Crucially, to mitigate biases in transcript quantification that can arise when highly expressed housekeeping genes dilute overall transcript per million (TPM) values within a genome, we implemented a dual quantification strategy. This included calculating BiG-MAP-derived TPMs^40^, which are normalized only against BGC-associated genes, enabling a more sensitive detection of BGC-specific induction (Fig.4a).

**Fig. 4.**
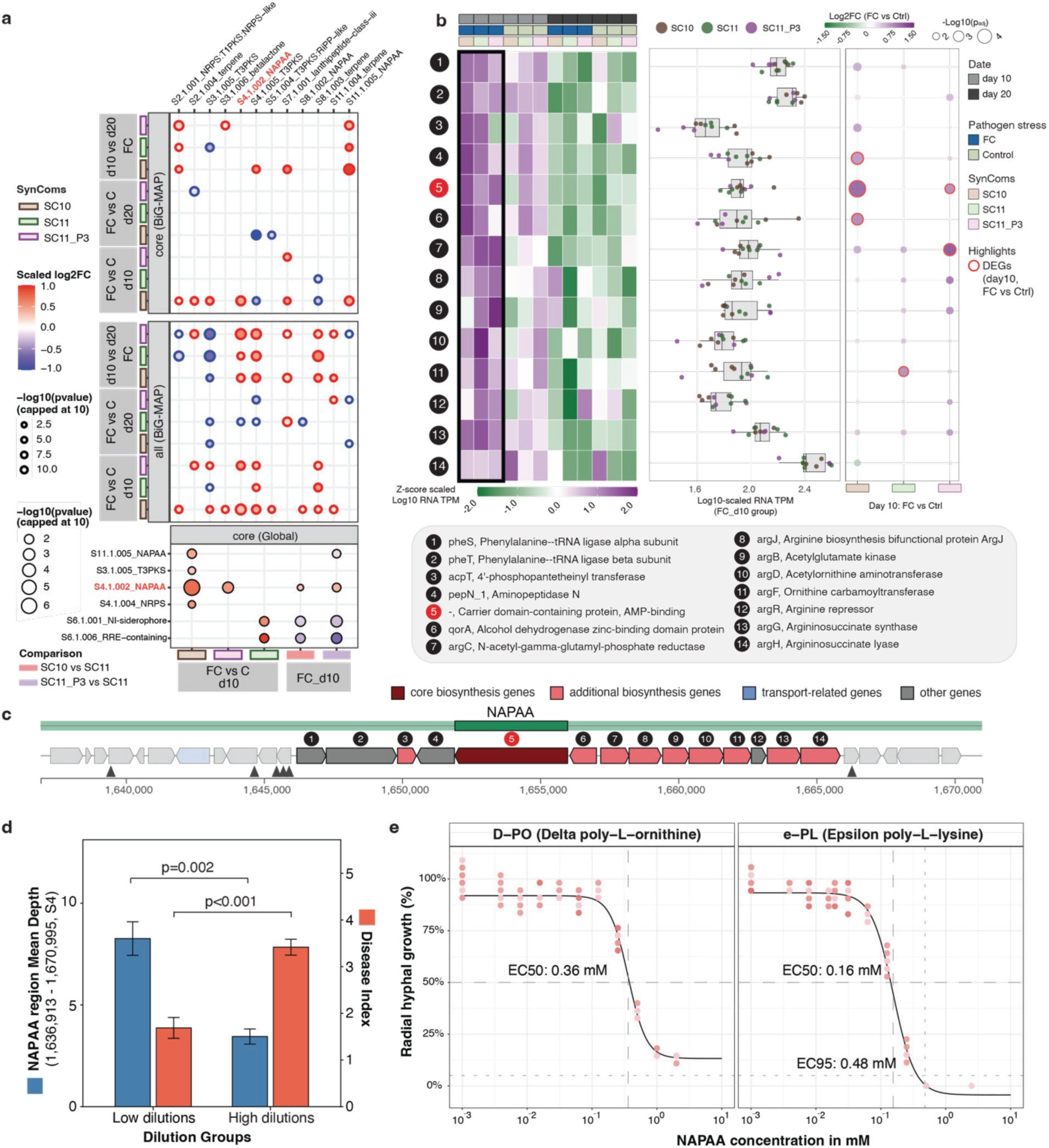
The biosynthetic gene clusters activated in the rhizosphere of disease-suppressive SynComs treated plants. **(a)** The heatmap dotplot shows the differentially expressed BGCs induced by pathogen or up-regulated in day 10. The dot colored by their log2-fold-change (Log2FC) value based on DESeq2 analysis using BiG-MAP normalized counts and original counts (Global). The BiG-MAP analysis provided normalized counts for both core region and full region (core, all). The global differential expression analysis was based on core genes of each BGC. The dot sizes are corresponded to −log10(p-adj-value) which indicate the significance between two groups. Color bar shows to which SynCom or comparison it belongs. **(b)** The heatmap illustrated the expression of NAPAA genes in all groups. The color bar on top of the heatmap shows the information of treatment including harvest date, pathogen stress, and SynCom treatment from top to down. The boxplot further zooms into the expression of *Fusarium*-inoculated day 10 group and displayed the RNA TPM in a Log10 value. The heatmap dotplot highlight the DEGs induced by pathogen at day 10 in red boarders. The dot color is based on Log2FC (DESeq2) and dot size indicate how significant the difference is. The chunk below the plots described the gene names and functions. **(c)** The structure of S4_NAPAA BGC. The length of each arrow corresponding to the gene length, and the color were coded by their antiSMASH gene functions (dark red: core biosynthetic genes; pink: additional biosynthetic genes; blue: transport related genes; green: regulatory gene; grey: other genes. Triangles: TTA codons). **(d)** Mean depth of NAPAA region in S4 strain and disease index across dilution groups according to DTE metagenome. Bar plots showing the mean depth (blue, left y-axis) and disease index (orange, right y-axis) for low dilution and high dilution treatment groups. Low dilution including 10X, 25X, 50X for disease index (36 samples), and 25X, 50X for mean depth (10 metagenome samples). High dilution including 100X, 200X, 400X for disease index (36 samples), and 100X, 200X for mean depth (10 metagenome samples). Bars represent group means; error bars indicate standard error of the mean. Statistical significance between dilution groups was assessed separately for each metric using the Mann–Whitney U test (p value in the plot). **(e)** *In vitro* growth inhibition of *Fusarium culmorum* by NAPAA compounds. Dose-response curves illustrate the inhibitory effects of two predicted NAPAA compounds, δ-poly-_L_-ornithine (D-PO) and ε-poly-_L_-lysine (e-PL), on the radial hyphal growth of *F. culmorum*. Assays were performed on water agar medium across a range of concentrations (10^-3^ to 2 mM). Individual biological replicates are represented by dots, with the color intensity (redness) of the points indicating a higher density of overlapping data points at that specific coordinate. The solid lines represent the best fit non-linear regression models used to determine the half-maximal effective concentration (EC50) and 95% reduction effective concentration (EC95). Data are normalized to the percentage of hyphal growth relative to the median value of untreated control group (100%).

Several BGCs, including betalactones, displayed elevated relative expression and differential regulation. In a complementary analysis, we correlated pathogen global transcriptional activity with the transcriptional activity of BGC-associated genes across samples and revealed significant positive associations in three NAPAA (non-alpha poly-amino-acid), two T3PKS, one terpene, and one lanthipeptide-class-III BGCs (Supplementary results; Fig.S22). Among these different analysis strategies, we consistently identified a NAPAA BGC from strain S4 and S6 that was significantly induced by the presence of the pathogen at day 10. The expression profile of this BGC across treatments demonstrated a highly coordinated induction specifically in the *Fusarium*-inoculated samples at the crucial day-10 time point (Fig.4b), establishing it as a strong candidate for a specialized metabolite at the onset of pathogen suppression.

The core gene of this novel cluster exhibited only 47% identity to the characterized epsilon-poly-_L_-lysine synthetase (BGC0002174), confirming its potential distinctiveness and suggesting it may encode either a typical polylysine or alternative non-alpha poly-amino-acid (NAPAA) product (Fig.S23). To assess the most likely products of this BGC, we then predicted the substrate specificity of the adenylation (A domain) using PARASECT, which indicated that lysine, ornithine, and arginine are the three likely substrates for the A domain (scores 0.633, 0.595, and 0.565, respectively). This substrate specificity range is further supported by the presence of a complete arginine biosynthetic operon (*argCJBDFRGH*)^41^ within the cluster vicinity (Fig.4b). Since ornithine is a known intermediate of this pathway and non-alpha polymerization of _L_-arginines is biosynthetically improbable, this suggests that δ-poly-_L_-ornithine is the most probable product of this novel NAPAA BGC. Furthermore, this BGC’s core gene was within the disease suppressiveness-related co-expression module *S4_M14* (Fig.S24). In this module, significant enrichment of genes related to amino acid biosynthesis, transport, and metabolism (highlighted in red color in Fig.S24) provides a clear transcriptional signature of a feed-forward mechanism, specifically assisting the NAPAA biosynthesis by provisioning the necessary amino acid precursors. The overall pathogen-induced pattern is supported by the identification of 23 differentially expressed genes (among 96 genes in this module) that were induced by the pathogen in at least one SynCom treatment. The core gene was confidently aligned with high identity (99.926%) to a contig (DTE_k127_28231982) from the DTE metagenome which did not assemble into any MAG. This DTE contig, in turn, was assigned to the S4 (*Arthrobacter equi*) isolate genome (99.863% identity). To validate the distribution of this BGC within the original DTE metagenome, reads were mapped to the NAPAA region (1,636,913 – 1,670,995). This analysis revealed that BGC abundance was significantly enriched in the low-dilution treatment (*p* = 0.002), correlating strongly with the observed levels of disease protection (*p* < 0.001) (Fig.4d).

To validate the antifungal potential of the NAPAA variants potentially encoded by this BGC, we had the most likely representative poly-amino acids chemically synthesized to perform *in vitro* growth inhibition assays against *F. culmorum*. We tested both ε-poly-_L_-lysine (ε-PL, microbial product of 25-35 residues) and δ-poly-_L_-ornithine (10 residues). Both compounds exhibited potent antifungal activity against *F. culmorum*, showing *EC50* values of 0.16 mM and 0.36 mM, respectively (Fig.4e). While the higher potency of ε-PL likely reflects its longer polymer chain, the activity of both variants confirms that NAPAA-type molecules are highly effective to suppress growth of *F. culmorum*.

## Discussion

In this study, we present a rationally designed and validated experimental and computational framework that successfully dissects a complex beneficial soil microbiome function to its mechanistic basis. Integrating dilution-to-extinction (DTE) perturbation with genome-resolved synthetic ecology overcame limitations inherent to low taxonomic resolution and ecological complexity, allowing us to isolate and functionally recapitulate a suppressive soil microbiome phenotype using an 11-member bacterial consortium.

The taxonomic composition of our 11-member SynCom (SynCom11) encompasses both well-known pathogen-suppressive bacterial genera (*Pseudomonas*^42,43^, *Arthrobacter*^29,44,45^, and *Microbacterium*^46,47^) and less commonly reported taxa (*Pedobacter*, *Mucilaginibacter*, *Amnibacterium*, and *Plantibacter cousiniae*), revealing diverse potential contributors to disease suppressiveness. *Pseudomonas* species have long been recognized as dominant plant beneficial bacteria through diverse mechanisms, especially the production of secondary metabolites that play key roles in the suppression of fungal diseases, such as 2,4-DAPG^34–37^, phenazines^14,48–50^, siderophores^51–54^, and cyclic lipopeptides^55–57^. Despite the presence of diverse secondary metabolite biosynthetic gene clusters, most *Pseudomonas* isolates from soil S11, including one strain (P3) that carried the 2,4-DAPG BGC, showed no significant negative correlation with disease severity in our DTE metagenomic analysis. To further interrogate the functional role of *Pseudomonas* in our system, we designed drop-out and replacement versions of SynCom11, alongside monoculture assays of the three *Pseudomonas* isolates (S1, P2, and P3). Remarkably, both SynCom11 and the *Pseudomonas*-free SynCom10 demonstrated robust disease suppression across replicated experiments, indicating that *Pseudomonas* members are most likely not critical for the overall disease-suppressive activity of soil S11. However, the replacement consortium (SynCom11_P3), in which the suppression-correlated strain was replaced by the 2,4-DAPG-harboring *Pseudomonas* strain, required sterile original S11 soil supplementation for consistent performance, and we observed phytotoxicity (leaf yellowing) in the SynCom11_P3 treatment under control conditions without pathogens. These findings suggest that *Pseudomonas*-mediated biocontrol is highly contingent on environmental context, and the presence of 2,4-DAPG biosynthetic potential does not inherently confer superior suppressive activity compared to other *Pseudomonas* strains. To test whether the consortium could be further simplified, two randomly selected half-depletion SynComs (containing only 5 and 6 strains, respectively) exhibited unstable disease suppression, suggesting cross-species interactions between multiple strains are critical for maintaining robust disease-suppressive activity.

Mechanistic investigations into plant disease suppression have often identified resource competition^12,58–60^, broad-spectrum antibiosis^43,61,62^, and chitinolytic activity^16,63,64^ as key modes of action. Our SynCom data critically challenges and expands these conventional strategies. The observed transcriptional activity of the T6SS aligns with established mechanisms of direct cellular antagonism and nutrient acquisition in the rhizosphere^65,66^. However, the expression of most siderophore BGCs was downregulated when the pathogen was present in the rhizosphere, which suggests that robust suppression was not due to iron competition. Expression analysis of chitinase-related genes showed that only a single chitinase gene was pathogen-induced, and this induction occurred solely in SynCom10, accompanied by considerable within-group variation across replicates. These findings suggest that chitinase activity is only one of the determinants of the disease-suppressive capacity observed for the SynComs. Functional analysis of the highly expressed genes revealed that many were associated with nitrogen utilization. Previous studies have demonstrated that microbial nitrogen transformation is strongly linked to disease-suppressive traits such as regulation of pathogen virulence^67^, beneficial bacterial colonization^68^, and limitation available nitrogen for pathogens^69^. Consistent with our DTE-based metagenomic results, a range of genes involved in amino acid synthesis, transport, and metabolism were pathogen-induced or transcriptionally upregulated. Notably, genes responsible for branched-chain amino acid biosynthesis were not only highly expressed at day 10 but also upregulated in the presence of the pathogen. This pattern was observed across multiple SynCom strains, suggesting that nitrogen flux and amino acid metabolism may play a crucial role in mediating community-level disease suppression.

Furthermore, the high transcriptional expression level of low-abundance *Arthrobacter equi* strains supported the assumption that rare taxa can function as disproportionately important drivers of ecosystem function^70^. Previous work on the genus *Arthrobacter* has highlighted its biocontrol and plant growth-promoting (PGP) capabilities through mechanisms like siderophore production, IAA synthesis, hydrolytic enzymes, and antagonistic metabolites^29,44,71,72^. Our findings uncovered the involvement of a novel non-alpha poly-amino-acid (NAPAA) BGC in soil suppressiveness. While NAPAA BGC is present across multiple SynCom members (S3, S4, S6, S7, S11), its pathogen-induced transcription and its integration into the dedicated co-expression module mark the low-abundance *Arthrobacter equi* (S4 and S6) as the functional linchpin for this strategy. The core biosynthesis gene of this cluster is predicted to encode an enzyme producing a NAPAA product, likely ε-poly-_L_-lysine, δ-poly-_L_-ornithine or (less chemically likely) ω-poly-_L_-arginine. Crucially, the biosynthesis pathway for these specific poly-amino acids remains largely uncharacterized, with recent work only reporting the δ-poly-_L_-ornithine pathway in *Acinetobacter baumannii*^73^. The finding of this novel NAPAA BGC and its pathogen-induced expression within the *Arthrobacter equi* strains is a major mechanistic advance, as no previous study has, to our knowledge, reported the contribution of any NAPAA BGC to plant-associated disease suppressiveness. The structural similarity to ε-poly-_L_-lysine (ε-PL), an approved safe food additive and preservative^74,75^, strongly suggests that this newly identified NAPAA class represents a promising source of safe, natural antimicrobial agents with significant potential for agricultural application. Although this study elucidates the functional significance of a novel NAPAA BGC in pathogen suppression, comprehensive characterization of its molecular structure, biocontrol mechanisms, and ecological distribution will require targeted approaches such as site-directed gene knockout analysis, heterologous expression systems, and extensive field validation.

Beyond the primary role of the NAPAA cluster, the correlation analysis between BGC transcription and fungal Relative Activity (RA) reveals a broader community-level response, and it is well possible that multiple parallel mechanisms are at play in a spatiotemporal manner. Specifically, within the SynComs, T3PKS clusters in strain S3 and S7 (*Plantibacter cousiniae*), as well as terpene module in S9 (*Amnibacterium*) and lanthipeptide modules in S10 (*Microbacterium hatanonis*), exhibited robust positive correlations (Spearman R > 0.8) with the pathogen RA (see more details in supplementary results). The co-activation of these distinct BGCs suggests that the SynCom may respond to pathogen-induced cues through a coordinated defense strategy. Additionally, while our workflow focuses on the cultivable bacterial fraction, we recognize that uncultured taxa identified as suppressive-correlated MAGs in our DTE analysis as well as other microbial taxa in the rhizosphere microbiome (e.g. fungi, algae, protists, viruses) may also contribute to the native soil’s complexity. Nevertheless, the fact that SynCom11 fully recapitulates the robust S11 suppressive phenotype demonstrates that our experimental and computational framework successfully distilled an essential functional core from the disease-suppressive microbiome.

Together, this work exemplifies how the systematic deconstruction of complex natural ecosystems can unlock the fundamental mechanisms that govern microbiome-mediated phenotypes. By successfully bridging top-down perturbations with bottom-up synthetic ecology, we provide a generalizable blueprint for transitioning from descriptive ‘omics’ to functional validation across diverse biomes. This framework, rooted in the integration of community-scale dynamics and strain-resolved molecular mechanisms, is readily adaptable to other complex terrestrial and aquatic systems, including plant, insect, animal and human microbiomes, where identifying the specific drivers of host health remains a central challenge. Ultimately, our approach offers a robust platform for the rational design of synthetic consortia and evidence-based biotechnological applications, paving the way for predictable microbiome engineering to address global challenges in agriculture, medicine, and environmental sustainability.

## Methods

### Soil and plant materials

Soil S11 was collected from an agricultural field near Bergen op Zoom, the Netherlands in March 2018. This soil exhibited a high level of disease suppressiveness against *Fusarium culmorum*, which was evaluated in our previous study^26^. Furthermore, we used a sandy dune soil (BS) collected near Bergharen, the Netherlands^76^. After collecting soils were air-dried in room temperature, sieved through 4 mm sieve, and stored at 4°C. BS soil was additionally gamma sterilized (Synergy Health Ede B.V., The Netherlands) before use.

### Seed preparation and growth conditions

JB Asano wheat seeds (Agrifirm, The Netherlands) were surface sterilized and pre-germinated in order to use in the experiment. Briefly, after surface sterilization with 70% ethanol for 1 minutes and 10% bleach for 10 minutes, seeds were washed with an excess of sterile water and placed on a wet 1mm paper filter (VWR, the Netherlands). Three-day-old seedlings were transferred to the soil, watered every second day and supplemented weekly with 0,5 Hoagland solution without microelements (0.5 M Ca(NO_3_)_2_·4H_2_O, 1 M KNO_3_, 1 M KH_2_PO_4_, 0.5 M MgSO_4_·7H_2_O and 98.6 mM ferric EDTA). Plants were grown in cabinets (MC 1750 VHO-EVD, Snijders Labs) at 20°C day and night, photoperiod 12 h day/12 h night with 60% relative humidity.

### Pathogen and microbiome inoculation and disease suppressive evaluation

Soilborne pathogen *F. culmorum* PV^77^was grown on 0.25 strength PDA media (Oxoid, The Netherlands) for 2 weeks at 20°C prior each experiment. Where indicated, the pathogen was introduced to the soil before seedling transfer as 6mm mycelium plugs (1 per 10cc of soil). After the experiment wheat plants were gently removed from the soil and cleaned. The disease symptoms on the roots were assessed according to a scale from 0- healthy to 5-severely diseased, like previously described^26^. Statistical differences in disease symptoms between treatments and control were assessed using the chi-square test, with an alpha cut-off of p<0.05.

### Microbiome extraction and dilution-to-extinction setup

Wheat plants were grown in four 130 cc pots filled with S11 for two weeks. In order to perform a liquid extraction of the microbiome from the suppressive soil S11, all the glassware, materials and buffer were autoclaved. Plants were gently removed from the pots and the soil with the whole root system was transferred to a glass bottle with 10 volumes of phosphate buffer (10 mM KH_2_PO_4_, pH=6.5) and 5 gram of sterile glass beads. These bottles were shaken using an orbital shaker for 1 h at 180 rpm, sonicated for 15 min and shaken again for 1h to detach microorganisms from soil particles. Afterwards, all the extracts were merged in one beaker and sieved through a metal sieve to separate bigger soil particles and roots. Subsequently, extract was briefly centrifuged (1000 x g, 1 min) and the supernatant was filtered again using a Buchner funnel through a filter paper (VWR Grade 415 Filter Paper, VWR the Netherlands). The extract obtained this way was considered a 10X diluted Extracted Soil Microbiome (ESM) because the soil was extracted with 10 volumes of the buffer. In parallel, as a control, the same procedure was applied with the sterile soil BS with a sterile vermiculite (Agra-vermiculite, The Netherlands) in a 1 to 1 volume ration. This way a BS control extract was obtained.

In order to optimize the magnitude of the dilution we performed preliminary experiments (data not shown) investigating first, if dilution affects soil disease suppressiveness to *F. culmorum* (the phenotype), and second, at which dilution point the change in phenotype from suppressive to conducive occurs. After the preliminary experiments we chose the dilution steps in such a way that the phenotype tipping point occurs centrally. Compared to the dilution steps used in other studies^22,78–80^, we applied a relatively small dilution magnitude. Nevertheless, in the experimental system we present in this work, these small dilution steps allow us to observe a gradual change of the phenotype.

The 10X dilution of the ESM was serially diluted with phosphate buffer to 25X, 50X, 100X, 200X and 400X and stored at 4°C. The dilution range was determined in a preliminary experiment (data not shown). Plastic, 100 cc pots were filled with the sterile soil BS mixed with sterile vermiculite in a volume ratio of 1 to 1 and with mycelium plugs of *F. culmorum* or sterile 0.25 PDA plugs (mock inoculation) according to the treatment. Distinct dilutions of the ESM and the BS control extract were added to pots, 20 ml per day for seven days, before wheat seedlings were introduced. Plants were grown for 3 weeks before the disease assessment and a rhizosphere DNA isolation.

All the dilution treatments and controls consisted of sixteen biological replicates; plants were grown in a randomized order to minimize spatial bias. A negative control and a positive control, with and without the pathogen, respectively, were used to control for the infectiousness of the pathogen and the effect of the sterile soil, buffer, and the extraction procedure.

The experiment was performed twice, and the rhizosphere samples used for the DNA extraction were collected in the second experiment from randomly selected replicates.

### Bacteria isolation, purification, and preservation

The rhizosphere bacteria were isolated from 10X dilution samples from the dilution-to-extinction experiment. Three replicate samples were pooled into one mixture, divided into 3 replicates (samples A, B, and C), and diluted into preferred dilutions before plating on different media. The 0.1 trypticase soy agar (TSA) medium, the King’s B (KB) agar medium, the VL-55 medium, and the chitin-yeast agar (CYA-chitin) medium were employed for the isolation of various taxa. A heat treatment was also performed at 80 °C for 20 min with sample B before plating for the isolation of heat-resistant bacteria. Plate 100 μL preferred dilutions (d1) as well as 10X (d2), and 100X (d3) into 5 plates per medium, incubated at 25 °C and start picking colonies after 3 days. The new colonies were picked every two days. each selected colony was streaked with a sterile toothpick on a new 0.1 TSA plate for purification. The purification step was repeated till the colonies were purified and the purified bacteria were stored with 40% glycerol at −80 °C with 4 replicates per strain.

### DNA extraction and whole genome sequencing for bacterial collections

The genomic DNA was extracted with bacterial pellets using a the Qiagen DNeasy PowerSoil Pro Kit protocol. The quality and quantity of the extracted DNA were assessed using a Qubit 4.0 Fluorometer and NanoDrop spectrophotometer. The genomes of these isolates were sequenced by Erasmus Medical Center with HiSeq2500 PE300. For strains involved in SynCom assay, we also performed sequencing with PromethION 2 Solo (Oxford Nanopore Technologies) by GenomeScan B.v. (https://genomescan.nl/).

### Synthetic community preparation, inoculation and experimental setup

The bioassay for screening disease-suppressive synthetic communities aimed to test the biocontrol potential of various synthetic communities, including monocultures of three *Pseudomonas* strains, against *Fusarium culmorum* in wheat plants. To prepare the pathogen inoculum, *F. culmorum* was cultured on 0.25 Potato Dextrose Agar (PDA) for 14 days at room temperature. Circular plugs (8 mm diameter) were excised from the culture using a sterile cork borer. Ten plugs were transferred to 15 mL Falcon tubes containing 10 mL of 0.5× Hoagland solution and mixed thoroughly.

For soil preparation, 50 mL of a sterile soil mixture (BS soil: vermiculite = 3:1, v/v) was transferred to an autoclaved beaker. The 10 mL of Hoagland solution containing *F. culmorum* plugs was added to the soil, mixed thoroughly with a sterile spoon, and then transferred to sterile 50 mL Falcon tubes. These tubes were incubated at 20°C for 8 days to allow pathogen establishment.

The synthetic communities (SynComs) were prepared by culturing each bacterial strains in 0.1× Tryptic Soy Broth (TSB) at 25°C with shaking at 180 rpm for 48 hours. The cultures were centrifuged, and the pellets were resuspended in 10 mM MgSO₄ solution. The optical density (OD) was measured at 600 nm, and the suspensions were adjusted to an OD of 0.02 for each member after mixing (i.e. final OD600 for 11-strain SynCom would be 0.22; for monoculture would be 0.02).

Surface-sterilized, pre-germinated wheat seedlings (2 days old) were treated with 5 mL of the SynCom solution (or 10 mM MgSO₄ solution for control groups) in petri dishes (5 seedlings per dish) and incubated at 20°C with gentle shaking (30 rpm) for 2 hours. The seedlings were then transferred to sterile petri dishes containing wet sterile filter papers and 10 mL of sterile demineralized water and incubated in a growth chamber at 20°C for 24 hours to facilitate bacterial colonization.

Subsequently, the pre-treated seedlings were transplanted into soil tubes that had been pre-incubated with *F. culmorum* for 8 days. The tubes were covered with Parafilm, leaving openings for plant growth, and were transferred to first 70% ethanol cleaned and afterwards UV-sterilized transparent boxes in the flow cabinet to prevent cross-contamination. Plants were grown in a walk-in growth chamber with a 12-hour light/12-hour dark cycle at 20°C. Sterile demineralized water was added every 3 days in flow cabinet, and a single supplementation of 0.5× Hoagland solution was provided on day 10. After 20 days, plants were carefully removed from the tubes, and their roots were washed with sterile water. The disease symptoms on the roots were assessed according at a scale from 0- healthy to 5-severely diseased, as described before.

To confirm the reproducibility of disease suppression and to harvest rhizosphere DNA, RNA, and metabolites, a similar bioassay was conducted with three selected disease-suppressive SynComs, incorporating some modifications. The schematic diagram of the SynCom-assay was shown in Fig.3a.

A sterile soil mixture (BS soil: S11 = 9:1, v/v) was used instead of the previous composition to improve DNA and RNA extraction quality. Pathogen inoculation was performed using *F. culmorum* spores (10⁵ spores per plant) suspended in 0.5× Hoagland solution. Additionally, pre-treated seedlings received an extra 1 mL of SynCom solution or 10 mM MgSO₄ solution (control) at transplantation to enhance colonization. Non-infected groups were included in this assay where the soil tubes were incubated for 14 days with 0.5× Hoagland solution without *F. culmorum* spores.

For this bioassay, 33 seedlings were treated and grown per experimental group. Fifteen plants were harvested on day 10, and the remaining 18 plants were harvested on day 20. Shoot length was measured at harvest, and shoot biomass were recorded after drying the tissues at 60°C for 5 days. Rhizosphere soil tightly associated with roots was collected in 50 mL Falcon tubes containing 10 mL of PowerProtect solution and stored at - 80°C for DNA and RNA extraction. Non-planted soil tubes were subjected to identical treatments and incubation to serve as blank samples for metabolite analysis. This approach ensured reliable assessment of disease suppression by SynComs while facilitating downstream molecular analyses.

### The chemical synthesis of delta poly-_L_-ornithine and antifungal test of NAPAA compounds

To evaluate the antifungal activity of the NAPAA BGC products, ε-poly-_L_-lysine was purchased commercially (CAS [28211-04-3]) and δ-poly-_L_-ornithine (10-mer) was chemically synthesized by LifeTein, LLC (https://www.lifetein.com/). Stock solution of these two compounds were prepared in sterile demi water and used to create a gradient of concentrations. These solutions were mixed into autoclaved water agar medium after it had cooled but before solidification. The amended media were then added into 6-well plates, with an equivalent volume of sterile water used for the control group. Following medium solidification, each well was centrally inoculated with 5 μL *Fusarium culmorum* spore suspension. Ten biological replicates were prepared for each treatment. Each 6-well plate contained one control and five distinct treatment concentrations. After 7 days incubation, the mycelia in the control group reached the edge of the well, the mycelial diameter was measured to quantify growth inhibition.

### Rhizosphere DNA and RNA extraction, library preparation and sequencing

In the DTE-assay, five out of sixteen replicates were randomly selected for a rhizosphere DNA extraction before the disease assessment. Plants were gently removed from the pots and the soil loosely adhering to the roots was removed by shaking. The whole root system was placed in a 50 ml falcon tube with 25 ml of sterile MQ grade water. The tubes were vortexed, sonicated, and vortexed again, each step for 1 min. Afterwards, the roots were removed with sterile forceps and disease symptoms were assessed on them. The water with extracted rhizosphere was briefly spined down (1000 x g, 1 min) and carefully decanted to a new 50 ml falcon tube. The decanted liquid was frozen at −20 °C and freeze dried overnight (Free zone 12, Labconco the USA). This resulted in forming a white powder in the tube which was collected and extracted using a DNeasy PowerSoil Kit (QIAGEN, the Netherlands) according to the manufacturer’s protocol, without applying a bead beating step. Samples were subsequently purified using the DNeasy PowerClean cleanup kit (QIAGEN, the Netherlands).

In the SynCom assay, the plants rhizosphere soil samples are collected in PowerProtect solution and stored at −20 degrees before the extraction. On the day before the DNA and RNA, the PowerProtect falcon tubes were moved to 4 degrees and defreeze overnight. The falcons were vortexed briefly and most of the soil particles were removed by pouring to a new 50 mL falcon tube. The PowerProtect solution was then completely removed by adding 20 mL sterile PBS buffer and centrifuge 9500 rcf for 10 min. The pellet was then resuspended with the PowerBeads solution provided in RNeasy PowerSoil Total RNA Kit (Qiagen) and transferred to the beads-tubes for the following RNA and DNA extraction. The RNA and DNA were extracted using RNeasy PowerSoil Total RNA Kit (Qiagen) and RNeasy PowerSoil DNA Elution Kit (Qiagen) follow the protocol provided with kits to have pairwise DNA and RNA samples. The obtained DNA and RNA were stored at −80 degrees before the library preparation, metagenomics and metatranscriptomics sequencing by Novogene b.v. (https://www.novogene.com/eu-en).

### Dilution-to-extinction (DTE) metagenome analyses

Raw paired sequence reads were quality-checked with fastqc (https://www.bioinformatics.babraham.ac.uk/projects/fastqc/) and trimmed using bbduk (Bushnell, n.d.) with a Phred score threshold of 30. Read pairs for which at least one mate was shorter than 150 bp were discarded. Reads for all dilution points were co-assembled in a single assembly using Megahit v1.2.9^81^. Genes were predicted using prodigal v2.6.3^82^ in metagenomic mode. Reads were mapped back onto the assembly contigs using hisat2^83^. BAM files were converted into SAM using samtools v1.7 and RPKM counts were obtained with mpileup^84^. Binning was performed within Anvi’o environment v6.2^85^ using DASTOOL^86^ refinement of bins obtained with MAXBIN2 v2.2.7^87^, CONCOCT v1.1.0^88^ and METABAT v2.15^89^. MAGs with a DASTOOL confidence score >= 0.5 were kept.

Identification of differentially abundant contigs was performed in R using DESeq v1.18.1^90^ looking for enrichment of contigs in the first two dilution points (25X and 50X) versus the higher dilution points (100X and 200X). Contigs with adjusted p-value < 0.05 were considered differentially abundant. Biosynthetic gene clusters were predicted from differentially abundant contigs with a positive fold change using antiSMASH 5.1.2^91^ using the prodigal-meta option for gene prediction, --cb-general and --cb-knownclusters for comparison of predicted clusters to the antiSMASH database^92^ and MIBiG^93^. Quality-trimmed reads were taxonomically annotated using Kraken v2.0.8^94^ against the GTDB-taxonomy database release 05-RS95^95^ KEGG annotation was performed at Baseclear (Leiden, The Netherlands) using the full megahit assembly.

Differentially abundant contigs which had at least one predicted cluster were taxonomically annotated using Diamond v 0.9.21^96^ against the NCBI nr database Oct. 2020^97^. Contigs were assigned to the taxonomical group with the highest cumulative score across the contig. Contigs annotated with the same taxonomy were grouped for tree construction. The list of taxonomic groups was used in NCBI common tree to obtain a tree in PHYLIP format representing the phylogeny relations of the taxonomies present in the differentially abundant contigs with biosynthetic gene clusters shown in Fig.S2a. Annotation of the tree was done with iTOL^98^; a fully annotated tree is available at https://itol.embl.de/tree/8873156232367181606086331#.

Presence-absence of KEGG orthologs enriched domains in MAGs was determined using hmmsearch v3.1b2^99^ and hmm domains from pfam^100^ against the different MAGs using trusted cutoffs. Presence of ClpV was determined based on the co-occurrence of all 4 ClpV subdomains (AAA, AAA_2, AAA_lid_9 and ClpB_D2-small) on the same contig. ClpV and Phage_base_V are considered core components of type VI secretion system^101^.

### Bacterial genome analyses and correlation analyses with DTE metagenome

The DNA were isolated from rhizosphere bacterial isolates mentioned in former section. The paired-end fastq files were trimmed by the fastp^102^, checked quality with the fastqc and multiQC^103^, short-reads genomes were assembled by the SPAdes^104^ and the long-reads genomes were assembled by the Flye^105^ and unicycler^106^. The assembled genomes were then checked for assembly quality with QUAST^107^, detected contamination with checkM2^108^, and decontaminated with MDMcleaner pipeline^109^. The genomes with low contamination and high completeness were then annotated with the functional genes using the eggNOG-mapper^110^, bakta^111^, and antiSMASH^91^. The biosynthetic gene clusters that identified with antiSMASH, were clustered into gene cluster families (GCFs) based on the similarities of the genes’ structure by BiG-SCAPE^112^. The replicated genomes were detected by fetchMGs^28^ with all marker genes with 100 % identity with each other. The phylogeny tree was constructed with GTDB-tk^113^, and the functional annotations were added to the tree using iTOL^98^.

The metagenomics raw reads of DTE-assay were aligned to assembled bacterial genomes separately using bowtie2^114^ and the statistical data were generated by samtools^84^. An R script was used to standardize the depth of mapped reads for each metagenome-genome pair, together with the phenotypic data, the correlation coefficient analysis was further performed to identify disease-related bacteria. The bacteria that were significantly negatively related to disease were selected for the SynComs’ assay. The DTE assembled contigs were mapped to both isolates genomes and MAGs using LexicMap^115^ with parameter settings: --min-qcov-per-genome = 60, --align-min-match-pident = 80, and --min-q-cov-per-hsp =60.

### Synthetic communities inoculated rhizosphere microbiome analyses

The raw reads of shotgun metagenomics and metatranscriptomics were first quality checked using fastQC and multiQC^103^, trimmed by fastp^102^, and then mapped to *Triticum aestivum* genome to remove host contamination using hisat2^83^. Then the clean reads were mapped to each isolated bacterial genomes using bowtie2^114^. The generated BAM files were then used to calculate the mapped counts for each genes using featureCounts^116^. The transcripts per million (TPM) were calculated for both metatranscriptome (defined as RNA TPM) and their paired metagenome (defined as DNA TPM), and the ratio of RNA_TPM/DNA_TPM was defined as TPM ratio. The calculation of TPM is:

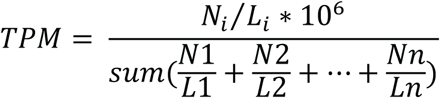

*N_i_* is the mapped counts for the *gene_i_*, *L_i_* is the length of the *gene_i_*, DNA TPM stands for using the metagenome raw reads mapped to each reference genomes to get the counts table, and RNA TPM was using metatranscriptome raw reads for mapping. The differential analysis for metatranscriptomics data was performed using DESeq2 and we used their paired metagenomics data as an offset (log (N_i_/N_total_)) to better select the differentially expressed genes/functions.

### Statistics and data visualizations

The statistical analysis and data visualization are all done with R. The R markdown files and original data using for the visualization are all provided in Zenodo: <u>10.5281/zenodo.18740830</u>.

## Data availability

Sequencing data for this study have been uploaded in the European Nucleotide Archive (ENA). The metagenome sequencing raw data of dilution-to-extinction are available under accession: PRJEB108303. The bacterial isolates assembled genomes are available under accession: PRJEB108483. The metagenomics and metatranscriptomics raw data of synthetic communities (SynComs) validation assay can be found under accession: PRJEB108647.

## Code availability

The codes for analysing can be downloaded from Zenodo: 10.5281/zenodo.18740830.

## Acknowledgements

This work was supported by the European Union via ERC Starting Grant 948770-DECIPHER to M.H.M., and the research program NWO-Groen, which is jointly funded by the Netherlands Organisation for Scientific Research (NWO), BASF SE, and Baseclear BV, under project number ALWGR.2015.1 (to M.H.M., G.P.v.W. and P.G.). J.J. was supported by China Scholarship Council (Grant No. 202106990013) for her PhD study.

## References

1. Olanrewaju, O. S., Glick, B. R. & Babalola, O. O. Beyond correlation: Understanding the causal link between microbiome and plant health. Heliyon 10, e40517 (2024).

2. Malard, F., Dore, J., Gaugler, B. & Mohty, M. Introduction to host microbiome symbiosis in health and disease. Mucosal Immunology 14, 547–554 (2021).

3. Banerjee, S. & Van Der Heijden, M. G. A. Soil microbiomes and one health. Nat Rev Microbiol 21, 6–20 (2023).

4. Singh, B. K., Yan, Z.-Z., Whittaker, M., Vargas, R. & Abdelfattah, A. Soil microbiomes must be explicitly included in One Health policy. Nat Microbiol 8, 1367–1372 (2023).

5. Raaijmakers, J. M. & Mazzola, M. Diversity and Natural Functions of Antibiotics Produced by Beneficial and Plant Pathogenic Bacteria. Annu. Rev. Phytopathol. 50, 403–424 (2012).

6. Gómez Expósito, R., De Bruijn, I., Postma, J. & Raaijmakers, J. M. Current Insights into the Role of Rhizosphere Bacteria in Disease Suppressive Soils. Front. Microbiol. 8, 2529 (2017).

7. Schlatter, D., Kinkel, L., Thomashow, L., Weller, D. & Paulitz, T. Disease Suppressive Soils: New Insights from the Soil Microbiome. Phytopathology® 107, 1284–1297 (2017).

8. Weller, D. M., Raaijmakers, J. M., Gardener, B. B. M. & Thomashow, L. S. Microbial populations responsible for specific soil suppressiveness to plant pathogens. Annu. Rev. Phytopathol. 40, 309–348 (2002).

9. Bakker, P. A. H. M. et al. The Soil-Borne Identity and Microbiome-Assisted Agriculture: Looking Back to the Future. Molecular Plant 13, 1394–1401 (2020).

10. Singh, B. K. et al. Climate change impacts on plant pathogens, food security and paths forward. Nat Rev Microbiol 21, 640–656 (2023).

11. Spooren, J. et al. Plant-Driven Assembly of Disease-Suppressive Soil Microbiomes. Annual Review of Phytopathology 62, 1–30 (2024).

12. Gu, S. et al. Competition for iron drives phytopathogen control by natural rhizosphere microbiomes. Nat Microbiol 5, 1002–1010 (2020).

13. Cha, J.-Y. et al. Microbial and biochemical basis of a Fusarium wilt-suppressive soil. The ISME Journal 10, 119–129 (2016).

14. Mazurier, S., Corberand, T., Lemanceau, P. & Raaijmakers, J. M. Phenazine antibiotics produced by fluorescent pseudomonads contribute to natural soil suppressiveness to Fusarium wilt. The ISME Journal 3, 977–991 (2009).

15. Berendsen, R. L. et al. Disease-induced assemblage of a plant-beneficial bacterial consortium. The ISME Journal 12, 1496–1507 (2018).

16. Carrión, V. J. et al. Pathogen-induced activation of disease-suppressive functions in the endophytic root microbiome. Science 366, 606–612 (2019).

17. Ossowicki, A., Raaijmakers, J. M. & Garbeva, P. Disentangling soil microbiome functions by perturbation. Environ Microbiol Rep 13, 582–590 (2021).

18. Siegel-Hertz, K. et al. Comparative Microbiome Analysis of a Fusarium Wilt Suppressive Soil and a Fusarium Wilt Conducive Soil From the Châteaurenard Region. Front. Microbiol. 9, 568 (2018).

19. Van Der Voort, M., Kempenaar, M., Van Driel, M., Raaijmakers, J. M. & Mendes, R. Impact of soil heat on reassembly of bacterial communities in the rhizosphere microbiome and plant disease suppression. Ecology Letters 19, 375–382 (2016).

20. Lee, S.-M., Kong, H. G., Song, G. C. & Ryu, C.-M. Disruption of Firmicutes and Actinobacteria abundance in tomato rhizosphere causes the incidence of bacterial wilt disease. The ISME Journal 15, 330–347 (2021).

21. Zegeye, E. K. et al. Selection, Succession, and Stabilization of Soil Microbial Consortia. mSystems 4, e00055–19 (2019).

22. Hol, W. H. G. et al. Non-random species loss in bacterial communities reduces antifungal volatile production. Ecology 96, 2042–2048 (2015).

23. Jing, J., Garbeva, P., Raaijmakers, J. M. & Medema, M. H. Strategies for tailoring functional microbial synthetic communities. The ISME Journal 18, wrae049 (2024).

24. Cambon, M. C., Xu, X., Kovács, Á. T., Gomes, S. I. & McDonald, J. E. Synthetic microbial communities for studying and engineering the tree microbiome: challenges and opportunities. Current Opinion in Microbiology 87, 102636 (2025).

25. Cairns, J. et al. Construction and Characterization of Synthetic Bacterial Community for Experimental Ecology and Evolution. Front. Genet. 9, 312 (2018).

26. Ossowicki, A. et al. Microbial and volatile profiling of soils suppressive to *Fusarium culmorum* of wheat. Proc. R. Soc. B. 287, 20192527 (2020).

27. Tracanna, V. et al. Dissecting Disease-Suppressive Rhizosphere Microbiomes by Functional Amplicon Sequencing and 10× Metagenomics. mSystems 6, e01116–20 (2021).

28. Milanese, A. et al. Microbial abundance, activity and population genomic profiling with mOTUs2. Nat Commun 10, 1014 (2019).

29. Yang, K. et al. Effects of Trichoderma harzianum and Arthrobacter ureafaciens on control of Fusarium crown rot and microbial communities in wheat root-zone soil. Phytopathol Res 7, 47 (2025).

30. Yin, C. et al. Role of Bacterial Communities in the Natural Suppression of Rhizoctonia solani Bare Patch Disease of Wheat (Triticum aestivum L.). Appl Environ Microbiol 79, 7428–7438 (2013).

31. Mavrodi, O. V., Walter, N., Elateek, S., Taylor, C. G. & Okubara, P. A. Suppression of Rhizoctonia and Pythium root rot of wheat by new strains of Pseudomonas. Biological Control 62, 93–102 (2012).

32. Tian, B. et al. Schizotrophic Sclerotinia sclerotiorum-Mediated Root and Rhizosphere Microbiome Alterations Activate Growth and Disease Resistance in Wheat. Microbiol Spectr 11, e00981–23 (2023).

33. Giannelli, G. et al. Exploring the rhizosphere of perennial wheat: potential for plant growth promotion and biocontrol applications. Sci Rep 14, 22792 (2024).

34. Raaijmakers, J. M. & Weller, D. M. Natural Plant Protection by 2,4-Diacetylphloroglucinol-Producing *Pseudomonas* spp. in Take-All Decline Soils. MPMI 11, 144–152 (1998).

35. Gong, L. et al. Novel synthesized 2, 4-DAPG analogues: antifungal activity, mechanism and toxicology. Sci Rep 6, 32266 (2016).

36. Weller, D. M. et al. Role of 2,4-Diacetylphloroglucinol-Producing Fluorescent *Pseudomonas* spp. in the Defense of Plant Roots. Plant Biology 9, 4–20 (2007).

37. Zhou, H. et al. Improving biocontrol activity ofPseudomonas fluorescens through chromosomal integration of 2,4-diacetylphloroglucinol biosynthesis genes. Chin.Sci.Bull. 50, 775–781 (2005).

38. Weissman, J. L., Hou, S. & Fuhrman, J. A. Estimating maximal microbial growth rates from cultures, metagenomes, and single cells via codon usage patterns. Proc. Natl. Acad. Sci. U.S.A. 118, e2016810118 (2021).

39. Choudhuri, B. S. et al. Overexpression and functional characterization of an ABC (ATP-binding cassette) transporter encoded by the genes drrA and drrB of Mycobacterium tuberculosis. Biochemical Journal 367, 279–285 (2002).

40. Pascal Andreu, V., et al. BiG-MAP: an Automated Pipeline To Profile Metabolic Gene Cluster Abundance and Expression in Microbiomes. mSystems 6, 10.1128/msystems.00937-21 (2021).

41. Duan, Y. et al. Fine-Tuning Multi-Gene Clusters via Well-Characterized Gene Expression Regulatory Elements: Case Study of the Arginine Synthesis Pathway in *C. glutamicum*. ACS Synth. Biol. 10, 38–48 (2021).

42. Mehmood, N. et al. Multifaceted Impacts of Plant-Beneficial *Pseudomonas* spp. in Managing Various Plant Diseases and Crop Yield Improvement. ACS Omega 8, 22296–22315 (2023).

43. Haas, D. & Keel, C. Regulation of antibiotic production in root-colonizing Pseudomonass spp. and relevance for biological control of plant disease. Annu. Rev. Phytopathol. 41, 117–153 (2003).

44. Ramlawi, S., Abusharkh, S., Carroll, A., McMullin, D. R. & Avis, T. J. Biological and chemical characterization of antimicrobial activity in *Arthrobacter* spp. isolated from disease-suppressive compost. J Basic Microbiol 61, 745–756 (2021).

45. Ramlawi, S., Aitken, A., Abusharkh, S., McMullin, D. R. & Avis, T. J. Arthropeptide A, an antifungal cyclic tetrapeptide from *Arthrobacter psychrophenolicus* isolated from disease suppressive compost. Natural Product Research 36, 5715–5723 (2022).

46. Ballot, A. et al. Dimethylpolysulfides production as the major mechanism behind wheat fungal pathogen biocontrol, by *Arthrobacter* and *Microbacterium* actinomycetes. Microbiol Spectr 11, e05292–22 (2023).

47. Patel, A. et al. New Insights on Endophytic Microbacterium-Assisted Blast Disease Suppression and Growth Promotion in Rice: Revelation by Polyphasic Functional Characterization and Transcriptomics. Microorganisms 11, 362 (2023).

48. Biessy, A. & Filion, M. Phenazines in plant-beneficial *Pseudomonas* spp.: biosynthesis, regulation, function and genomics. Environmental Microbiology 20, 3905–3917 (2018).

49. Chin-A-Woeng, T. F. C., Bloemberg, G. V. & Lugtenberg, B. J. J. Phenazines and their role in biocontrol by *Pseudomonas* bacteria. New Phytologist 157, 503–523 (2003).

50. Mavrodi, D. V., Blankenfeldt, W. & Thomashow, L. S. Phenazine Compounds in Fluorescent *Pseudomonas* Spp. Biosynthesis and Regulation. Annu. Rev. Phytopathol. 44, 417–445 (2006).

51. Feng, Z. et al. A synthetic community of siderophore-producing bacteria increases soil selenium bioavailability and plant uptake through regulation of the soil microbiome. Science of The Total Environment 871, 162076 (2023).

52. Sasirekha, B. & Srividya, S. Siderophore production by Pseudomonas aeruginosa FP6, a biocontrol strain for Rhizoctonia solani and Colletotrichum gloeosporioides causing diseases in chilli. Agriculture and Natural Resources 50, 250–256 (2016).

53. Sayyed, R. Z., Chincholkar, S. B., Reddy, M. S., Gangurde, N. S. & Patel, P. R. Siderophore Producing PGPR for Crop Nutrition and Phytopathogen Suppression. in Bacteria in Agrobiology: Disease Management (ed. Maheshwari, D. K.) 449–471 (Springer Berlin Heidelberg, Berlin, Heidelberg, 2013). doi:10.1007/978-3-642-33639-3_17.

54. Shao, Z. et al. Siderophore interactions drive the ability of *Pseudomonas* spp. consortia to protect tomato against Ralstonia solanacearum. Horticulture Research 11, uhae186 (2024).

55. De Souza, J. T., De Boer, M., De Waard, P., Van Beek, T. A. & Raaijmakers, J. M. Biochemical, Genetic, and Zoosporicidal Properties of Cyclic Lipopeptide Surfactants Produced by *Pseudomonas fluorescens*. Appl Environ Microbiol 69, 7161–7172 (2003).

56. Gu, Y., Ma, Y., Wang, J., Xia, Z. & Wei, H. Genomic insights into a plant growth-promoting *Pseudomonas koreensis* strain with cyclic lipopeptide-mediated antifungal activity. MicrobiologyOpen 9, e1092 (2020).

57. Omoboye, O. O. et al. Pseudomonas Cyclic Lipopeptides Suppress the Rice Blast Fungus Magnaporthe oryzae by Induced Resistance and Direct Antagonism. Front. Plant Sci. 10, 901 (2019).

58. Kinkel, L. L., Schlatter, D. C., Bakker, M. G. & Arenz, B. E. Streptomyces competition and co-evolution in relation to plant disease suppression. Research in Microbiology 163, 490–499 (2012).

59. Sharp, R. T., Shaw, M. W. & Van Den Bosch, F. The effect of competition on the control of invading plant pathogens. Journal of Applied Ecology 57, 1403–1412 (2020).

60. Dutt, A., Andrivon, D. & Le May, C. Multi-infections, competitive interactions, and pathogen coexistence. Plant Pathology 71, 5–22 (2022).

61. Wang, X. Q., Zhao, D. L., Shen, L. L., Jing, C. L. & Zhang, C. S. Application and Mechanisms of Bacillus subtilis in Biological Control of Plant Disease. in Role of Rhizospheric Microbes in Soil (ed. Meena, V. S.) 225–250 (Springer Singapore, Singapore, 2018). doi:10.1007/978-981-10-8402-7_9.

62. Arseneault, T. & Filion, M. Biocontrol through antibiosis: exploring the role played by subinhibitory concentrations of antibiotics in soil and their impact on plant pathogens. Canadian Journal of Plant Pathology 39, 267–274 (2017).

63. Zhang, X. et al. Enhancing plant disease suppression by Burkholderia vietnamiensis through chromosomal integration of Bacillus subtilis chitinase gene chi113. Biotechnol Lett 34, 287–293 (2012).

64. Zhang, Z., Yuen, G. Y., Sarath, G. & Penheiter, A. R. Chitinases from the Plant Disease Biocontrol Agent, *Stenotrophomonas maltophilia* C3. Phytopathology® 91, 204–211 (2001).

65. Yin, R., Cheng, J. & Lin, J. The role of the type VI secretion system in the stress resistance of plant-associated bacteria. Stress Biology 4, 16 (2024).

66. Song, L. et al. Trojan horselike T6SS effector TepC mediates both interference competition and exploitative competition. The ISME Journal 18, wrad028 (2024).

67. Guan, X. et al. Microbial nitrogen transformation regulates pathogenic virulence in soil environment. Journal of Environmental Management 369, 122280 (2024).

68. Sós-Hegedűs, A. et al. Suppression of *NB-LRR* genes by miRNAs promotes nitrogen-fixing nodule development in *MEDICAGO TRUNCATULA*. Plant Cell & Environment 43, 1117–1129 (2020).

69. Snoeijers, S. S., Pérez-García, A., Joosten, M. H. A. J. & De Wit, P. J. G. M. The Effect of Nitrogen on Disease Development and Gene Expression in Bacterial and Fungal Plant Pathogens. European Journal of Plant Pathology 106, 493–506 (2000).

70. Chen, Q.-L. et al. Rare microbial taxa as the major drivers of ecosystem multifunctionality in long-term fertilized soils. Soil Biology and Biochemistry 141, 107686 (2020).

71. Chhetri, G. et al. An Isolated Arthrobacter sp. Enhances Rice (Oryza sativa L.) Plant Growth. Microorganisms 10, 1187 (2022).

72. Gonçalves, O. S. et al. Harnessing Novel Soil Bacteria for Beneficial Interactions with Soybean. Microorganisms 11, 300 (2023).

73. Patel, K. D. & Gulick, A. M. Structural and functional insights into δ-poly-L-ornithine polymer biosynthesis from Acinetobacter baumannii. Commun Biol 6, 982 (2023).

74. Hyldgaard, M. et al. The Antimicrobial Mechanism of Action of Epsilon-Poly- L -Lysine. Appl Environ Microbiol 80, 7758–7770 (2014).

75. Rodrigues, B. et al. Antimicrobial activity of Epsilon-Poly-l-lysine against phytopathogenic bacteria. Sci Rep 10, 11324 (2020).

76. Schulz-Bohm, K., Zweers, H., De Boer, W. & Garbeva, P. A fragrant neighborhood: volatile mediated bacterial interactions in soil. Front. Microbiol. 6, (2015).

77. Chen, Q.-L., Ding, J., Zhu, Y.-G., He, J.-Z. & Hu, H.-W. Soil bacterial taxonomic diversity is critical to maintaining the plant productivity. Environment International 140, 105766 (2020).

78. Korenblum, E. et al. Rhizosphere microbiome mediates systemic root metabolite exudation by root-to-root signaling. Proc. Natl. Acad. Sci. U.S.A. 117, 3874–3883 (2020).

79. Yan, Y., Kuramae, E. E., Klinkhamer, P. G. L. & Van Veen, J. A. Revisiting the Dilution Procedure Used To Manipulate Microbial Biodiversity in Terrestrial Systems. Appl Environ Microbiol 81, 4246–4252 (2015).

80. Li, D. et al. MEGAHIT v1.0: A fast and scalable metagenome assembler driven by advanced methodologies and community practices. Methods 102, 3–11 (2016).

81. Hyatt, D. et al. Prodigal: prokaryotic gene recognition and translation initiation site identification. BMC Bioinformatics 11, 119 (2010).

82. Kim, D., Paggi, J. M., Park, C., Bennett, C. & Salzberg, S. L. Graph-based genome alignment and genotyping with HISAT2 and HISAT-genotype. Nat Biotechnol 37, 907–915 (2019).

83. Li, H. et al. The Sequence Alignment/Map format and SAMtools. Bioinformatics 25, 2078–2079 (2009).

84. Eren, A. M. et al. Anvi’o: an advanced analysis and visualization platform for ‘omics data. PeerJ 3, e1319 (2015).

85. Sieber, C. M. K. et al. Recovery of genomes from metagenomes via a dereplication, aggregation and scoring strategy. Nat Microbiol 3, 836–843 (2018).

86. Wu, Y.-W., Simmons, B. A. & Singer, S. W. MaxBin 2.0: an automated binning algorithm to recover genomes from multiple metagenomic datasets. Bioinformatics 32, 605–607 (2016).

87. Qian, J. & Comin, M. MetaCon: unsupervised clustering of metagenomic contigs with probabilistic k-mers statistics and coverage. BMC Bioinformatics 20, 367 (2019).

88. Kang, D. D. et al. MetaBAT 2: an adaptive binning algorithm for robust and efficient genome reconstruction from metagenome assemblies. PeerJ 7, e7359 (2019).

89. Love, M. I., Huber, W. & Anders, S. Moderated estimation of fold change and dispersion for RNA-seq data with DESeq2. Genome Biol 15, 550 (2014).

90. Blin, K. et al. antiSMASH 5.0: updates to the secondary metabolite genome mining pipeline. Nucleic Acids Research 47, W81–W87 (2019).

91. Blin, K., Shaw, S., Kautsar, S. A., Medema, M. H. & Weber, T. The antiSMASH database version 3: increased taxonomic coverage and new query features for modular enzymes. Nucleic Acids Research 49, D639–D643 (2021).

92. Kautsar, S. A. et al. MIBiG 2.0: a repository for biosynthetic gene clusters of known function. Nucleic Acids Research gkz882 (2019) doi:10.1093/nar/gkz882.

93. Wood, D. E., Lu, J. & Langmead, B. Improved metagenomic analysis with Kraken 2. Genome Biol 20, 257 (2019).

94. Parks, D. H. et al. A complete domain-to-species taxonomy for Bacteria and Archaea. Nat Biotechnol 38, 1079–1086 (2020).

95. Buchfink, B., Xie, C. & Huson, D. H. Fast and sensitive protein alignment using DIAMOND. Nat Methods 12, 59–60 (2015).

96. Database resources of the National Center for Biotechnology Information. Nucleic Acids Res 44, D7–D19 (2016).

97. Letunic, I. & Bork, P. Interactive Tree Of Life (iTOL) v4: recent updates and new developments. Nucleic Acids Research 47, W256–W259 (2019).

98. Mistry, J., Finn, R. D., Eddy, S. R., Bateman, A. & Punta, M. Challenges in homology search: HMMER3 and convergent evolution of coiled-coil regions. Nucleic Acids Research 41, e121–e121 (2013).

99. El-Gebali, S. et al. The Pfam protein families database in 2019. Nucleic Acids Research 47, D427–D432 (2019).

100. Boyer, F., Fichant, G., Berthod, J., Vandenbrouck, Y. & Attree, I. Dissecting the bacterial type VI secretion system by a genome wide in silico analysis: what can be learned from available microbial genomic resources? BMC Genomics 10, 104 (2009).

101. Chen, S., Zhou, Y., Chen, Y. & Gu, J. fastp: an ultra-fast all-in-one FASTQ preprocessor. Bioinformatics 34, i884–i890 (2018).

102. Ewels, P., Magnusson, M., Lundin, S. & Käller, M. MultiQC: summarize analysis results for multiple tools and samples in a single report. Bioinformatics 32, 3047–3048 (2016).

103. Bankevich, A. et al. SPAdes: A New Genome Assembly Algorithm and Its Applications to Single-Cell Sequencing. Journal of Computational Biology 19, 455–477 (2012).

104. Kolmogorov, M., Yuan, J., Lin, Y. & Pevzner, P. A. Assembly of long, error-prone reads using repeat graphs. Nat Biotechnol 37, 540–546 (2019).

105. Wick, R. R., Judd, L. M., Gorrie, C. L. & Holt, K. E. Unicycler: Resolving bacterial genome assemblies from short and long sequencing reads. PLoS Comput Biol 13, e1005595 (2017).

106. Gurevich, A., Saveliev, V., Vyahhi, N. & Tesler, G. QUAST: quality assessment tool for genome assemblies. Bioinformatics 29, 1072–1075 (2013).

107. Chklovski, A., Parks, D. H., Woodcroft, B. J. & Tyson, G. W. CheckM2: a rapid, scalable and accurate tool for assessing microbial genome quality using machine learning. Nat Methods 20, 1203–1212 (2023).

108. Vollmers, J., Wiegand, S., Lenk, F. & Kaster, A.-K. How clear is our current view on microbial dark matter? (Re-)assessing public MAG & SAG datasets with MDMcleaner. Nucleic Acids Research 50, e76–e76 (2022).

109. Cantalapiedra, C. P., Hernández-Plaza, A., Letunic, I., Bork, P. & Huerta-Cepas, J. eggNOG-mapper v2: Functional Annotation, Orthology Assignments, and Domain Prediction at the Metagenomic Scale. Molecular Biology and Evolution 38, 5825–5829 (2021).

110. Schwengers, O. et al. Bakta: rapid and standardized annotation of bacterial genomes via alignment-free sequence identification: Find out more about Bakta, the motivation, challenges and applications, here. Microbial Genomics 7, (2021).

111. Navarro-Muñoz, J. C. et al. A computational framework to explore large-scale biosynthetic diversity. Nat Chem Biol 16, 60–68 (2020).

112. Sunagawa, S. et al. Metagenomic species profiling using universal phylogenetic marker genes. Nat Methods 10, 1196–1199 (2013).

113. Chaumeil, P.-A., Mussig, A. J., Hugenholtz, P. & Parks, D. H. GTDB-Tk: a toolkit to classify genomes with the Genome Taxonomy Database. Bioinformatics 36, 1925–1927 (2020).

114. Langmead, B. & Salzberg, S. L. Fast gapped-read alignment with Bowtie 2. Nat Methods 9, 357–359 (2012).

115. Shen, W., Lees, J. A. & Iqbal, Z. Efficient sequence alignment against millions of prokaryotic genomes with LexicMap. Nat Biotechnol https://doi.org/10.1038/s41587-025-02812-8 (2025) doi:10.1038/s41587-025-02812-8.

116. Liao, Y., Smyth, G. K. & Shi, W. featureCounts: an efficient general purpose program for assigning sequence reads to genomic features. Bioinformatics 30, 923–930 (2014).

