## Supplementary information for "Reconstructing plant beneficial bacterial consortia by integrating dilution-to-extinction microbiome perturbation with genome-resolved synthetic ecology"

### Supplementary Materials for

The file includes:

Supplementary Results

Figs. S1 to S24

References for supplementary results

#### Supplementary Results

*Genes, biosynthetic gene clusters (BGCs) and metagenome-assembled genomes associated with suppression from DTE metagenome analysis*

Co-assembly of all rhizosphere metagenome reads generated 327,947 contigs (> 5 Kb, N50 = 11.9 Kb), which we analyzed to identify potential suppression-associated genomic features. We identified 42,170 differentially abundant contigs between suppressive and conducive samples, with 504 BGCs assigned to these contigs (Fig.S2a). BGC analysis revealed enrichment of melanin, aryl polyene, and T3PKS biosynthesis pathways in suppressive samples (Table S1), while functional annotation identified 147 enriched KEGG orthologs (KOs) (Table S2), including genes associated with iron acquisition like TonB [ko: K03832], bacterioferritin [ko: K03594], chitinases [ko: K01183], and components of the Type VI Secretion System (T6SS) [ko: K11894]. Further integration of these features into Metagenome-Assembled Genomes (MAGs) identified nine specific bins, including *Burkholderiaceae*, *Acidimicrobiaceae*, *Verrucomicrobiaceae*, and *Polyangiales*, that were strongly associated with suppressiveness and co-harbored these prioritized functions (Supplementary Results; Fig.S2b, c). Although siderophore biosynthesis has been observed as enriched function in our previous work[1] and has been linked to soil suppressiveness against *Fusarium oxysporum* in other studies[2–4], metallophore BGC abundance in our samples did not vary significantly with the loss of suppressiveness. Chitinases have been reported as a primary mechanism of endophytic bacteria conferring soil suppressiveness in sugar beet against *Rhizoctonia solani*[5]. Likewise, T6SS are tightly associated with antagonism of bacteria against pathogen through the secretion of toxins and pathogenicity factors[6–10].

Given the extensive number of potential targets contigs and KOs, we further prioritized the ones that are assigned to differential metagenome-assembled genomes (MAGs) along the DTE trajectory to provide additional genomic and taxonomic context. Manual curation of individual bins resulted in a collection of 96 MAGs, with 9 of them strongly associated with disease suppressiveness (Fig. S2b) and they all contain at least two of the three prioritized functions (iron-competition-, chitinase-, and T6SS- related functions)

(Fig. S2c). These suppression-associated MAGs belonged to the bacterial families *Burkholderiaceae*, *Acidimicrobiaceae*, *Verrucomicrobiaceae*, and *Polyangiales*. Thus, the top-down DTE approach revealed specific bacterial families and functional genes potentially associated with disease suppressiveness, but highlighted the need for complementary culture-based approaches to expand the limited resolution of the MAG-based analysis. For the BGCs encoding melanins, aryl polyenes, and ladderanes (Table.S1), melanins have previously been implicated in rhizosphere colonization[11], but little evidence supports their direct role in disease suppressiveness; the same holds for aryl polyenes and ladderanes.

*Link between metagenome analysis and suppressive SynComs*

The overall *in situ* expression analysis provided insights for further analysis, and we next validated the restoration and expression of the prioritized genes (selected via DTE metagenome analysis) within the SynCom-treated rhizosphere focusing on day 10. Specifically, we present an integrated analysis across four distinct analysis layers (Fig.S16a-e) focused on: (i) suppressive-associated KEGG Orthology (KO) identified from the initial DTE metagenome (orange); (ii) KOs demonstrating high gene expression levels (purple, RNA TPM > 100 in the day10_FC groups); (iii) KOs induced by pathogen at day 10 (blue) and (iv) KOs exhibiting heightened activation at day 10, coinciding with highest pathogen-bacterial interactions (green). The results show that across the annotated KOs in SynCom genomes (3892 KOs), about 31% (1208 KOs) were upregulated at day 10, 18% (693 KOs) were induced by pathogen, and about 12% (476 KOs) were transcriptionally highly expressed. Among DTE metagenome selected KOs (Table.S2), 121 KOs were restored with the SynComs, around 21% (25 KOs) were with high gene expression level (purple), and 31% (37 KOs) were induced by pathogen (Fig.S16b&d). The top 50 expressed DTE-prioritized-KOs in *Fusarium*-inoculated day 10 samples (Fig.S16c) revealed that the biosynthesis of secondary metabolites and ribosome pathways were highly expressed. Notably, type IV pili (T4P) were also activated in suppressive-SynCom rhizosphere (Fig.S16e).

We further analysed the expression of iron-uptake, T6SS, and chitinase related genes and found that only T6SS and ferritin were upregulated (Fig.S16f). Most of the siderophore BGCs as well as the TonB genes closed to these BGCs were downregulated in the presence of the pathogen (Fig.S17). In total 331 chitinase related genes (K01183, K01207, K01233, K01452) were found in the SynCom member genomes, but only one gene (S6_18975_K01183) was highly expressed (median RNA TPM > 100) (Fig.S12). This indicate that iron-competition and chitinase may not be the essential mechanisms for the suppression brought by the SynComs. The T6SS pathway was completely highly activated in S5 strain which indicates a potential contribution to the suppression phenotype. Beyond the DTE prioritized functions, we also revealed that the branched-chain amino acid (BCAAs) biosynthesis pathway was present in all other 3 analysis layers (Fig.S18). The complete biosynthesis pathway of BCAAs exhibited coordinated upregulation, with pathogen-induced expression detected across multiple strains (S2, S5, S10, S11) (Fig.S18).

*Highly transcriptionally expressed genes in Fusarium-inoculated rhizosphere microbiome*

Lollipop visualization of the highly expressed genes in *Fusarium*-inoculated treatments further revealed that highly active gene predominantly originated from seven community members (*Mucilaginibacter*, *Arthrobacter equi*, *Pedobacter*, *Plantibacter cousiniae*, *Microbacterium*, *Amnibacterium*), especially *Arthrobacter equi* S4 and S6 (Fig.S9). These genes were linked to nitrogen metabolism, stress response (i.e. *cspA*[12], *aldA*[13]), and biofilm formation, with stress response genes being crucial for bacterial survival during microbe-pathogen interactions. Nitrogen metabolism stood out, with key genes like *amt*[14] ( ammonium transporter, import $NH_4^+$), *mauE*[15] (methylamine utilization protein, process organic nitrogen), *thiS*[16] (thiamine biosynthesis carrier, cofactor for nitrogen-related enzymes), and *nirD*[17] (nitrite reductase subunit). Radar charts profiling nitrogen metabolism gene (Fig.S19) confirmed that *Arthrobacter equi* (S4 and S6) encoded the most nitrogen metabolism genes, all of which were highly expressed and pathogen-induced in SynCom 10 and SynCom 11. *Plantibacter cousiniae* (S3 and S7) also contained many activated nitrogen metabolism genes, though fewer pathogen-induced. The *Pseudomonas* strains (S1, P3), showed a highly activated biofilm formation function (Fig.S20). The enrichment of nitrogen metabolism genes suggests that efficient nitrogen acquisition and cycling may be fundamental to the biocontrol capacity, potentially through enhanced nutrient competition with pathogens[18,19] or improved plant nitrogen nutrition[20].

Other potential BGCs that may contribute to disease suppression

From the correlation analysis, we identified three NAPAA, two T3PKS BGCs, one terpene, and one lanthipeptide-class-iii BGCs have significant positive associations with disease severity. S3_BGC1.8 and S7_BGC1.4 (*Plantibacter cousiniae*) are both 17% identity compared to BGC0000934 from *Kitasatospora sp. HKI 741*; S4_BGC1.2, S6_BGC1.2 (*Arthrobacter equi*), S11_BGC1.5 (*Microbacterium kitamiense*) were identified to BGC0002174 (ε-Poly-L-lysine from *Epichloe festucae*); S9_BGC61.1 is aligned with 9% to BGC0002504 (nocardiopsistin A from *Nocardiopsis sp.*); and S10_BGC1.1 (*Microbacterium hatanonis*, lanthipeptide-class-iii) is not aligned to any known BGC, but there are example of lanthipeptide compound that have antifungal activity[21]. The T3PKS related BGCs were often present in biocontrol agent like *Pseudomonas*[22,23], *Streptomyces*[24,25], and *Bacillus*[26–28]. The terpene BGCs were linked more to *Nocardiopsis*[29,30] and little evidence regarding to their biocontrol potential in *Amnibacterium*.

**Fig. S1 to S24**

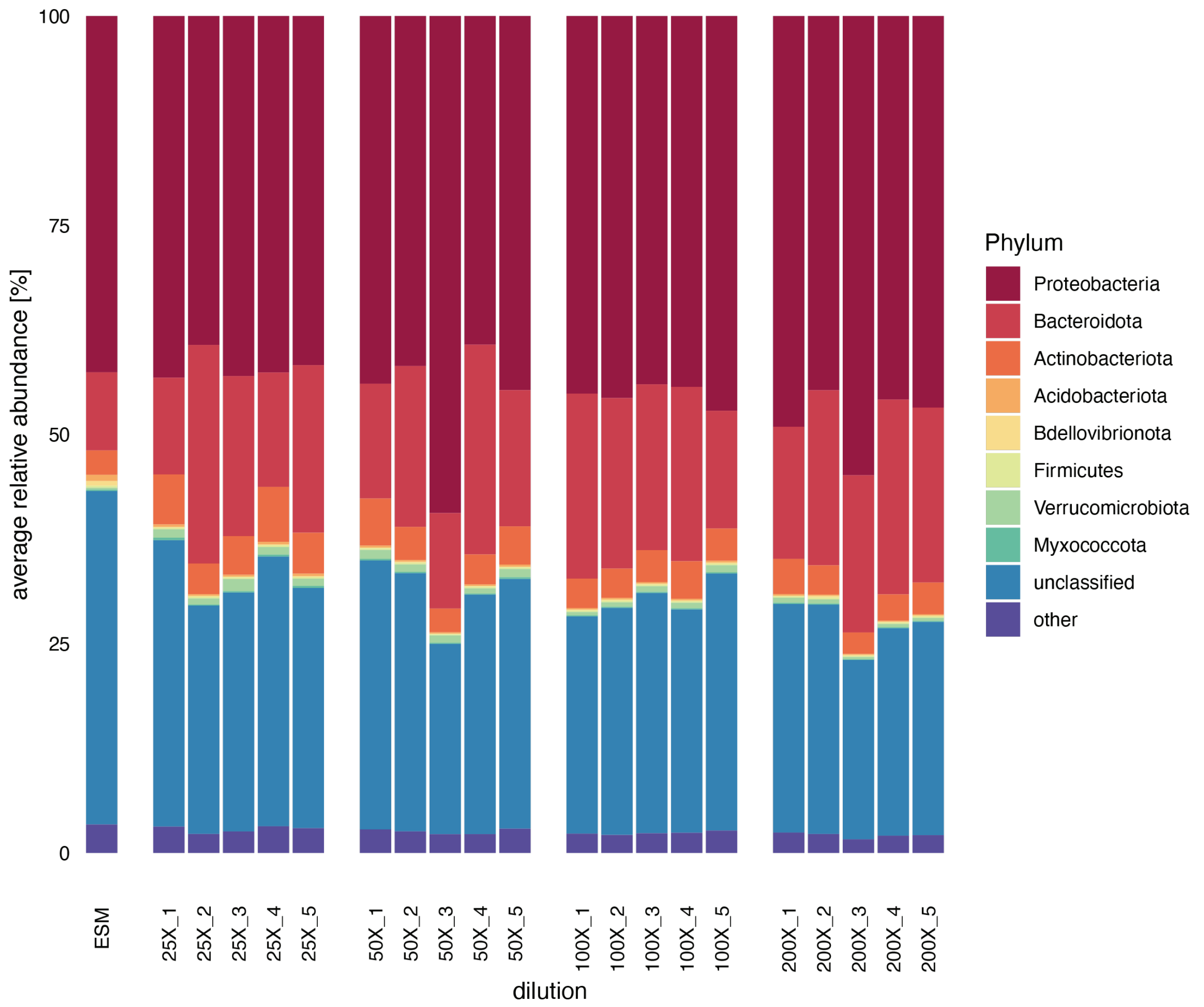


**Fig.S1. Barplot showing the most abundant phyla in the ESM and rhizosphere across four dilution points in five replicates.** Different color represents different phylum.

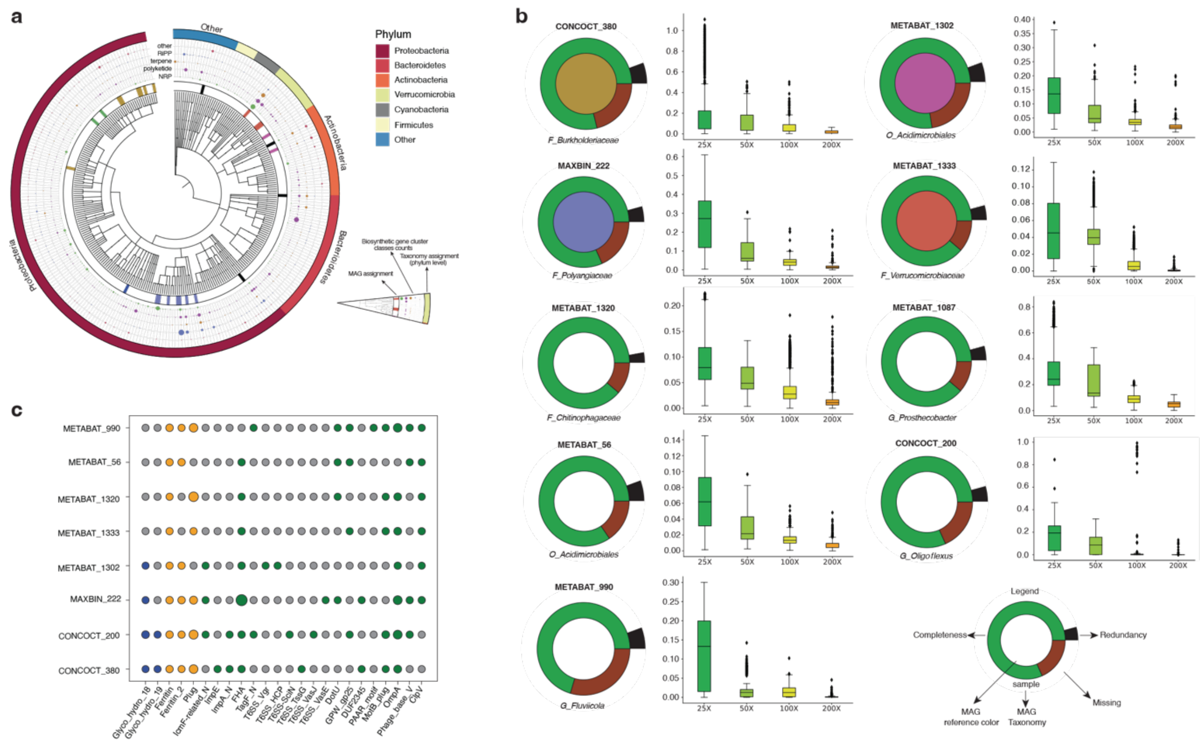

**Fig.S2. Taxonomical and functional representation of 504 differential abundant contigs containing BGCs from the dilution metagenomes and 9 suppression-associated MAGs. (a)** A phylogenetic tree based on their taxonomical annotation is used to order the contigs. In this visulization, contigs assigned to the same taxa are grouped together. The innermost circle (MAG assignment) represents the taxonomic groups containing contigs assigned to a specific metagenome-assembled genome. Contigs assigned to the same MAG are marked with the same color or with black for MAGs containing only one BGC-harboring contig. Further, the major antiSMASH-predicted BGC classes in the contigs are indicated with dots. The number of predicted BGC is reflected by the size of the dot. Finally, the outer ring represents the contig taxonomical assignment at the phylum level. **(b)** Representation of the MAGs from the metagenome showing significant association with the suppressive phenotype. The MAGs shown here are constituted by 50% or more in length by differentially abundant contigs according to DESeq2 analysis. The ring shows an overview of the MAGs assembly completeness and redundancy. For each MAG, a boxplot shows the abundance of all its contigs across the dilutions. MAG reference colors refer to panel a. The taxonomic levels are indicated by a capital letters: O - Order, F - Family, and G - genus. **(c)** Presence (colored dot) absent (gray dot) patterns of domains contained in KEGG orthologues genes enriched in suppressive dilution points in the MAG collection. Bacterial chitinase families are shown in blue, siderophores associated domains in orange and components of type VI secretion system (T6SS) in green. ClpV is considered present (last column) if all 4 ClpV subdomain (AAA, AAA_2, AAA_lid_9 and ClpB_D2-small) cooccur on the same contig.

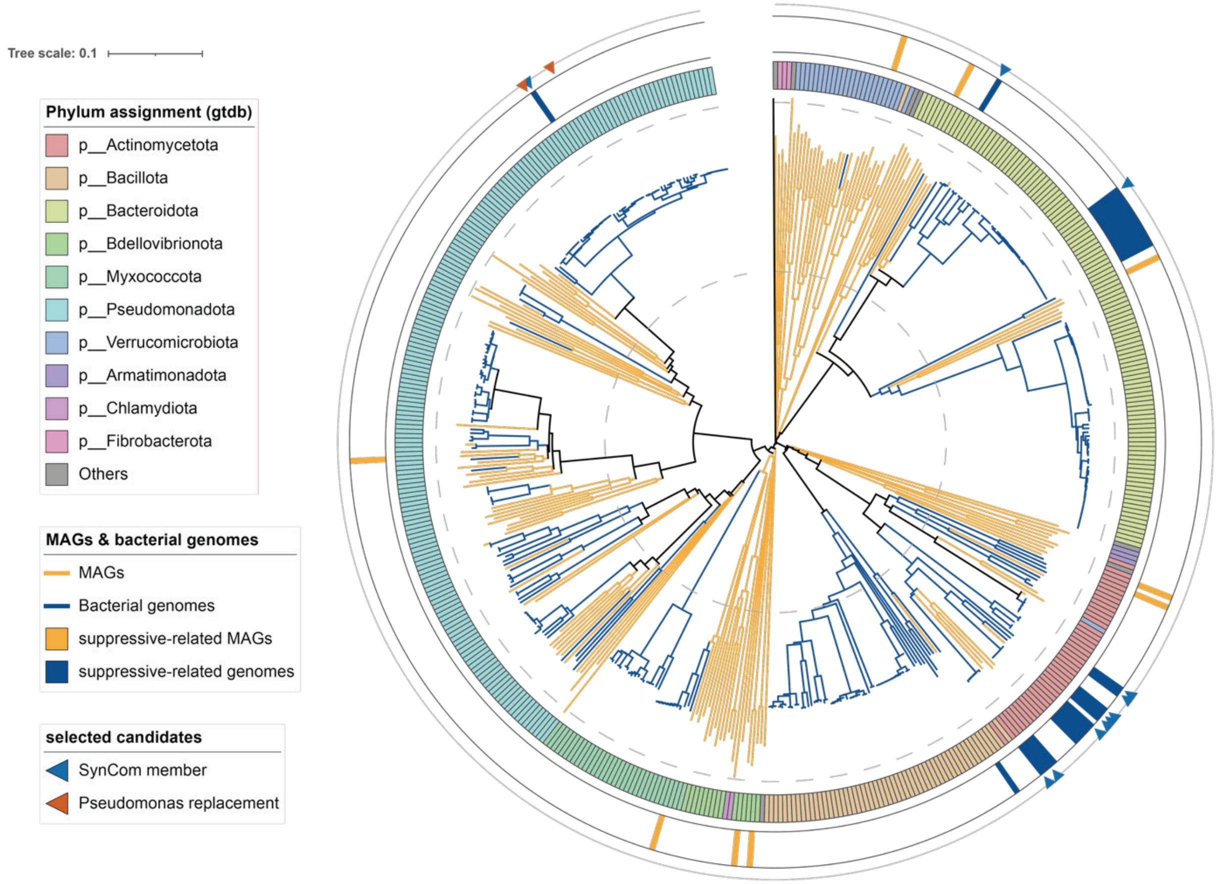


**Fig.S3. The comparison between DTE metagenome-assembled genomes (MAGs) and bacterial isolates' genomes.** The branch colors indicate the genomes resource, orange color represent MAGs and blue stand for bacterial genomes. The inner cycle color bar shows the phylum assignment according to gtdb database. The median cycle highlighted the MAGs and the genomes that are significantly negatively associated with disease index. The outer cycle highlighted the candidates that were selected for SynComs test.

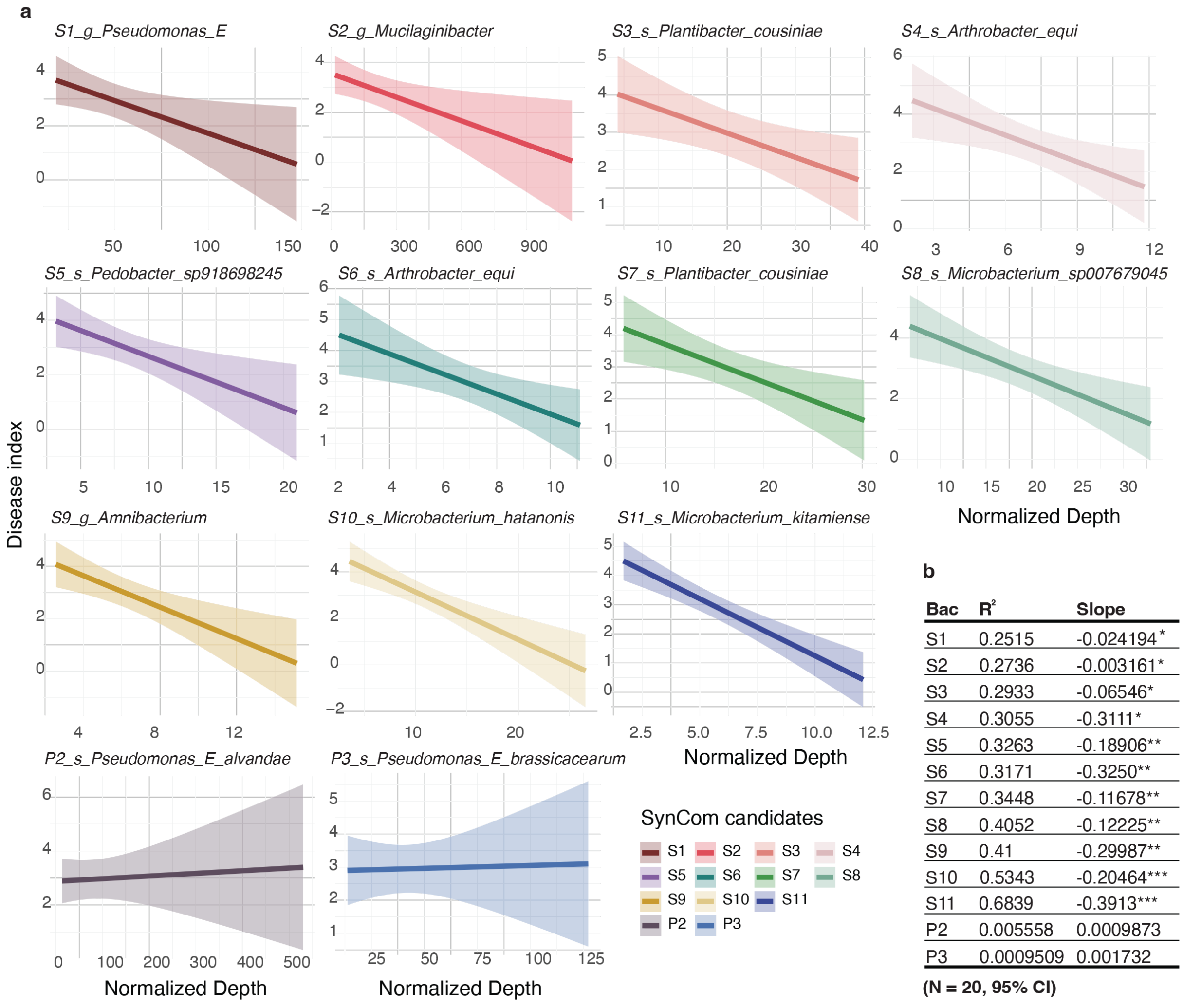


**b**

| Bac | $R^2$ | Slope |
|---|---|---|
| S1 | 0.2515 | -0.024194* |
| S2 | 0.2736 | -0.003161* |
| S3 | 0.2933 | -0.06546* |
| S4 | 0.3055 | -0.3111* |
| S5 | 0.3263 | -0.18906** |
| S6 | 0.3171 | -0.3250** |
| S7 | 0.3448 | -0.11678** |
| S8 | 0.4052 | -0.12225** |
| S9 | 0.41 | -0.29987** |
| S10 | 0.5343 | -0.20464*** |
| S11 | 0.6839 | -0.3913*** |
| P2 | 0.005558 | 0.0009873 |
| P3 | 0.0009509 | 0.001732 |

**(N = 20, 95% CI)**

**Fig.S4. The linear correlation between disease index and the abundance of SynCom candidates**. **(a)** The linear model plots of each SynCom candidate. The x-axis is the normalized depth of each candidate in DTE samples, and the y-axis is the disease index (0-5) of that sample. Shaded areas represent 95% confidence intervals. Colors indicate different bacterial isolates as shown in the legend. **(b)** Summary table showing the regression slope and coefficient of determination ($R^2$) for each strain. Negative slopes indicate that higher abundance of the corresponding strain is associated with reduced disease severity.

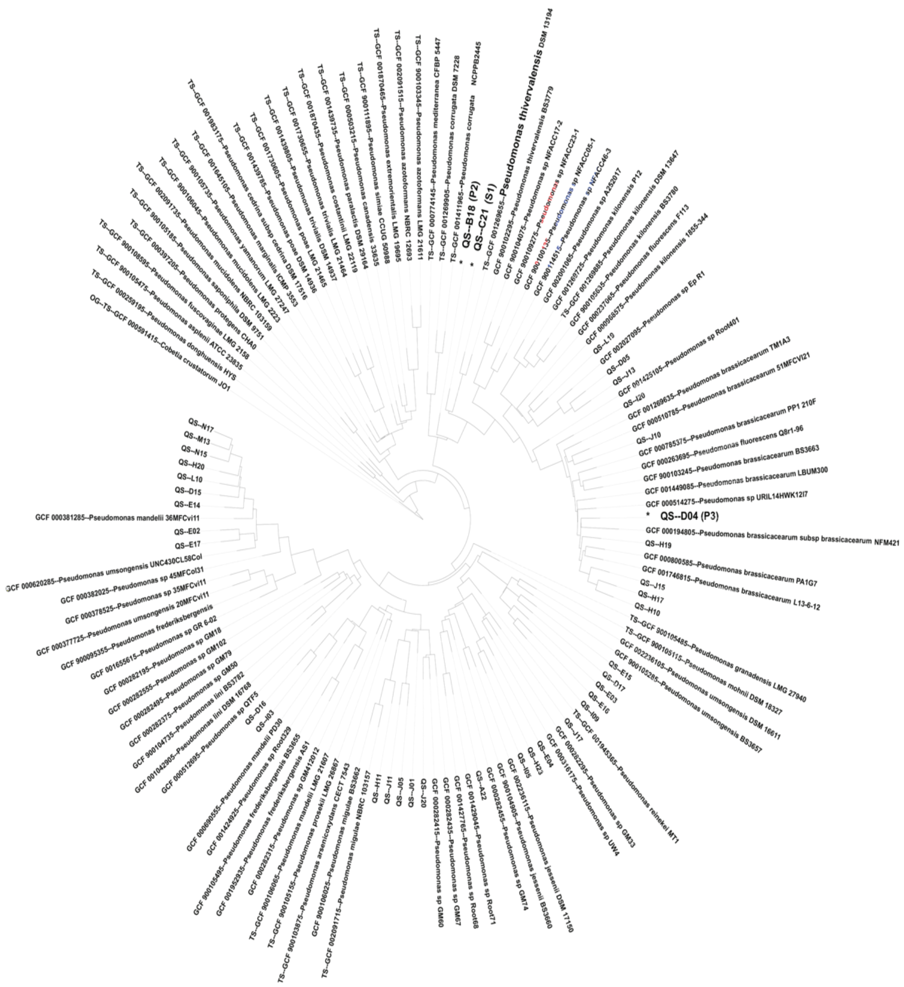


**Fig.S5. A phylogenetic tree of *Pseudomonas* isolates and closest reference genomes.** The blue color highlight *Pseudomonas* S1 strain which has linear negative correlation with disease index. The red color highlights the *Pseudomonas* P2 (the closest strain next to S1) and P3 (the 2,4-DAPG BGC containing strain) strains that selected to be tested in the SynCom.

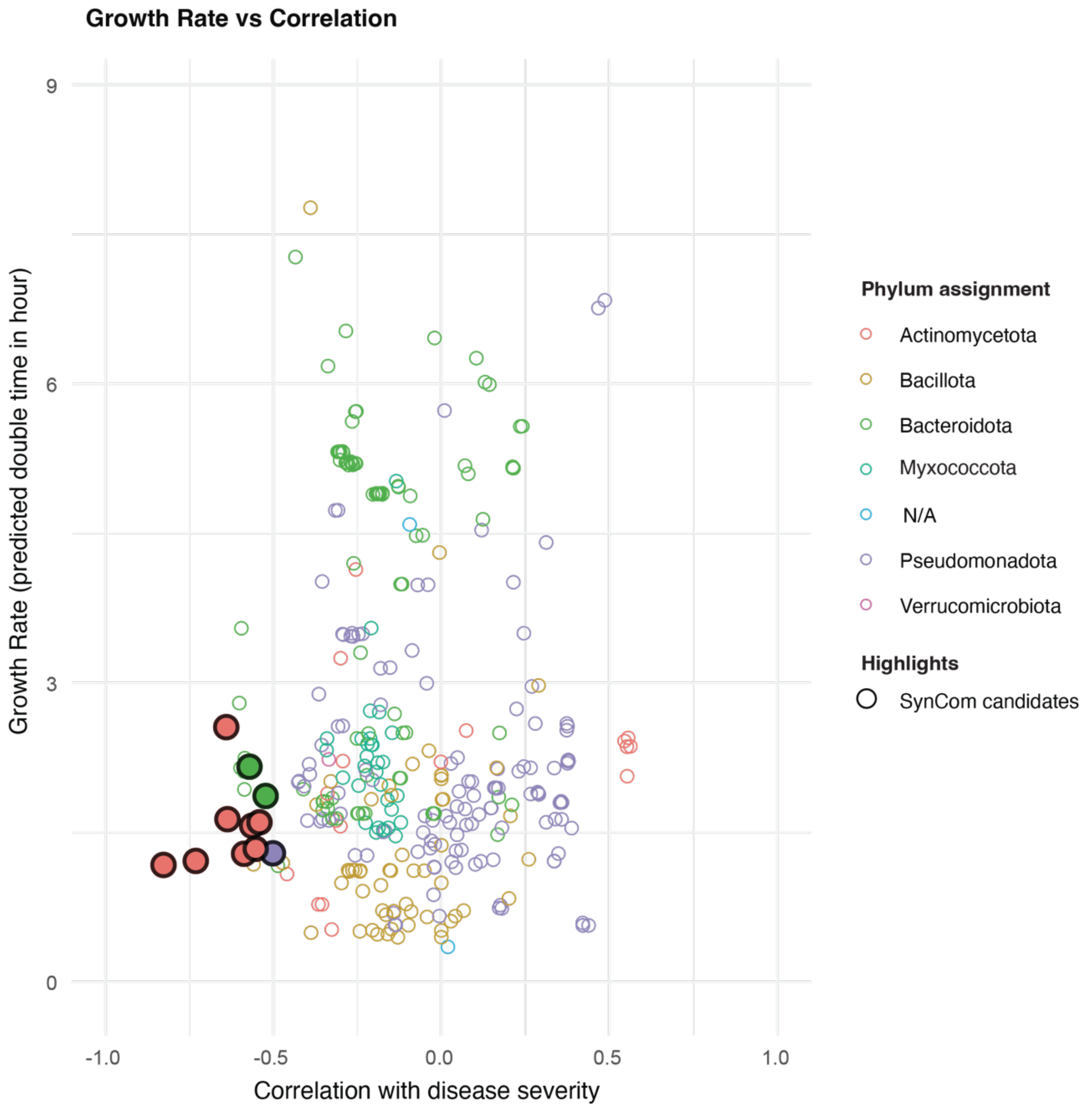


**Fig.S6. Growth rate versus correlation with disease severity for bacterial isolates.** Each point represents an isolate, with the x-axis showing the correlation of its relative abundance with disease severity and y-axis indicating the predicted doubling time (hours). Boarder colors correspond to phylum-level taxonomic assignments. Candidate strains selected for synthetic community (SynCom) are highlighted with larger colored dots.

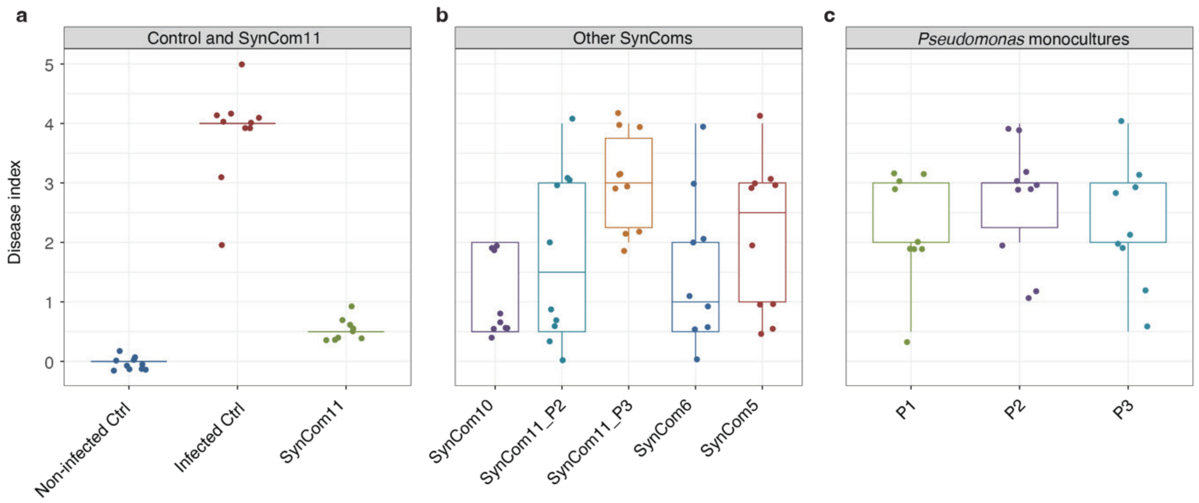


**Fig.S7. A complete plant phenotype outcome from SynCom experiment 1.** Disease severity was evaluated across three experimental conditions. **(a)** Controls infected and non-infected plants and SynCom11 treatment. **(b)** Alternative SynCom combinations. SynCom10: SynCom11 without *Pseudomonas* S1 strain; SynCom11_P2: replace S1 to P2 from SynCom11; SynCom11_P3: replace S1 to P3 from SynCom11; SynCom6: combination of S1, S3, S5, S7, S9, and S11; SynCom5: combination of S2, S4, S6, S8, and S10. **(c)** Individual *Pseudomonas* monocultures. P1 (also named S1): *Pseaudomonas* strain that show negative correlation to disease index in DTE experiment. P2: *Pseudomonas* strain closely related to P1 but no correlation with disease index in DTE experiment. P3: *Pseudomonas* strain with 2,4-DAPG BGC. Each point represents an individual replicate, horizontal lines indicate medians, and boxplots show interquartile ranges with whiskers extending to 1.5x the interquartile range.

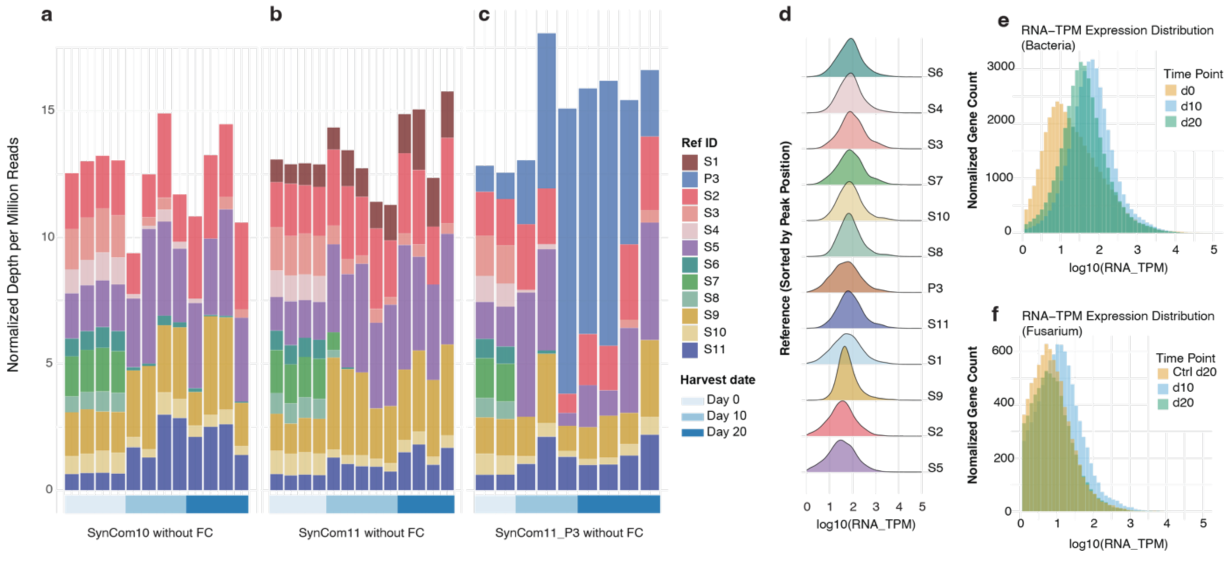


**Fig.S8. The changes in SynCom composition without *Fusarium culmorum* and global transcriptional activity in different samples. (a-c)** The barplots show the normalized depth of the inoculated bacteria in the rhizosphere of 3 SynComs without pathogen stress based on shotgun metagenomics data. Color-bar from light to dark blue represents

DNA samples collected at day 0, day 10, and day 20, respectively. Each color indicates a different bacterial member of SynCom. Day 0 samples represent the initial inoculum, while day 10 and day 20 samples were collected from rhizosphere soil. The barplot was evaluated across three SynComs without pathogen infection. **(a)** SynCom10 **(b)** SynCom11 **(c)** SynCom11_P3. **(d)** The ridge plot displays the expression distribution of inoculated bacteria in the *Fusarium*-inoculated groups at day 10. Reference genomes are arranged from top to bottom in descending order of peak expression levels. **(e-f)** The ridge plot displays the expression distribution of inoculated SynComs and *Fusarium culmorum* in different date. Gene counts were normalized by samples size for each group. For comparing *Fusarium culmorum* activity with/without SynComs, samples collected from day 20 were split into "Ctrl d20" (non-SynCom-inoculated plants) and "d20" (SynCom-inoculated plants).

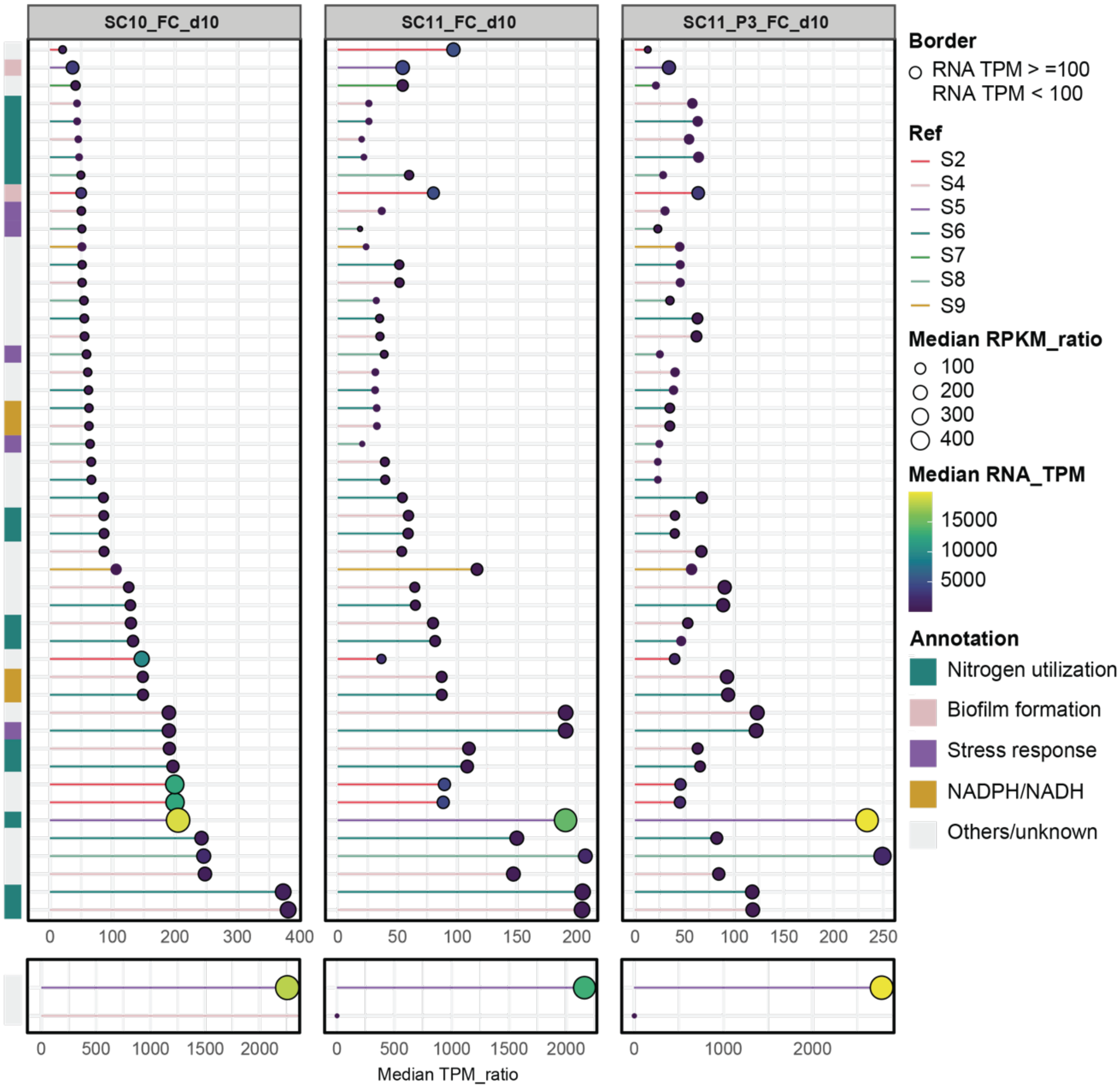


**Fig.S9. Top expressed genes in suppressive synthetic communities.** The lollipop plot illustrated the highly expressed genes in *Fusarium*-inoculated treatments from day 10. Only genes with a TPM ratio (RNA TPM / DNA TPM) greater than 50 are shown. Bar lengths represent the median TPM ratio across biological replicates. Bar color represents their reference genome. Circles size indicates the median RPKM ratio (RNA RPKM / DNA RPKM). Circles outlines distinguish transcriptional activity for which RNA TPM ≥ 100 with black boarder, < 100 with red boarder. Circles color shows the median RNA TPM value according to the adjacent color scale. Color bar next to the plot shows their functional assignment.

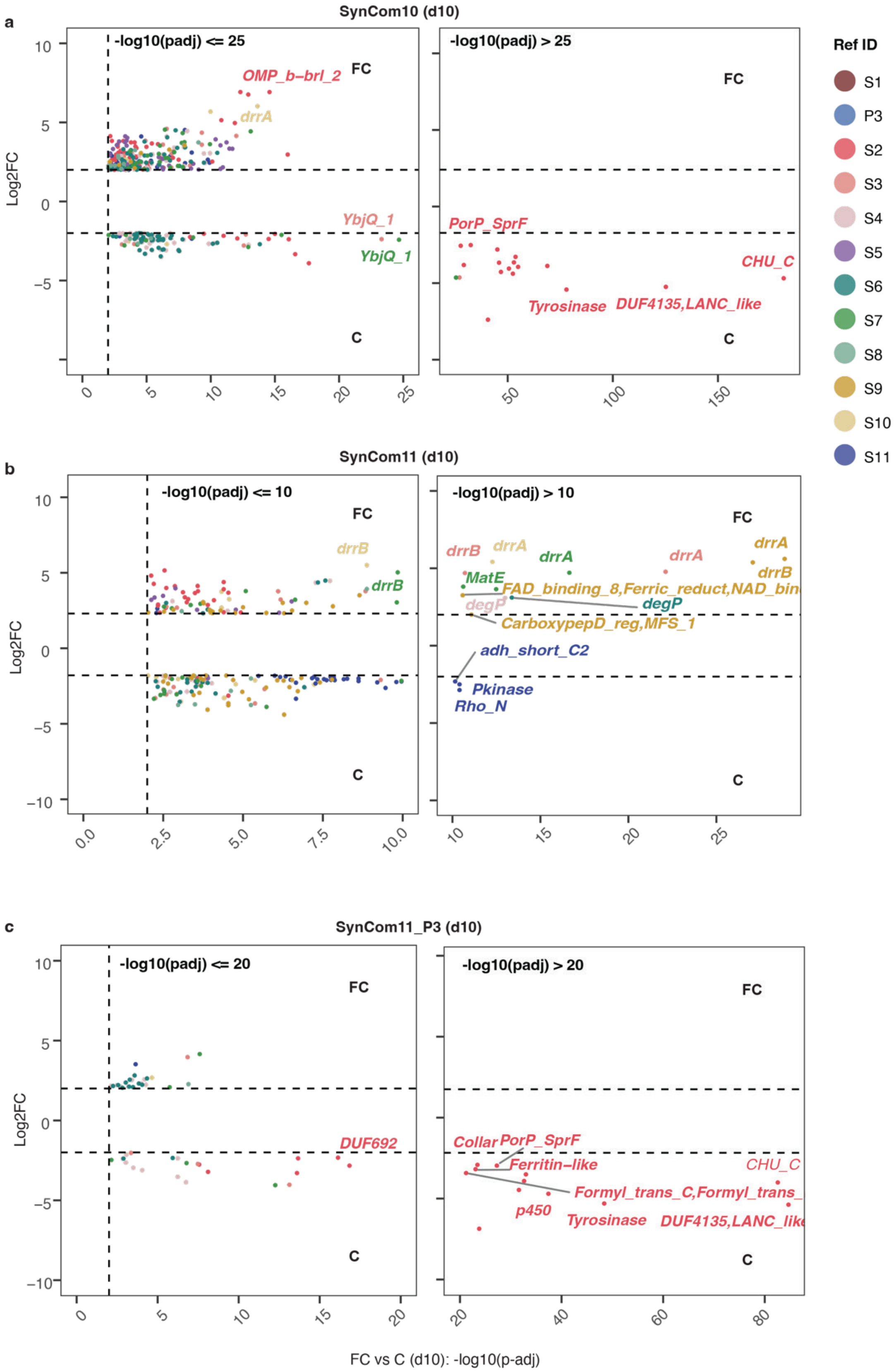
a
SynCom10 (d10)
-log10(padj) <= 25
-log10(padj) > 25
Log2FC
FC
C
OMP_b-brl_2
drrA
YbjQ_1
YbjQ_1
PorP_SprF
CHU_C
Tyrosinase
DUF4135,LANC_like
Ref ID
S1
P3
S2
S3
S4
S5
S6
S7
S8
S9
S10
S11
b
SynCom11 (d10)
-log10(padj) <= 10
-log10(padj) > 10
drrB
drrB
drrB
drrA
drrA
drrA
drrA
drrB
MatE
FAD_binding_8,Ferric_reduct,NAD_bin
degP
degP
CarboxypepD_reg,MFS_1
adh_short_C2
Pkinase
Rho_N
c
SynCom11_P3 (d10)
-log10(padj) <= 20
-log10(padj) > 20
DUF692
Collar
PorP_SprF
Ferritin-like
CHU_C
Formyl_trans_C,Formyl_trans_
p450
Tyrosinase
DUF4135,LANC_like
FC vs C (d10): -log10(p-adj)

**Fig.S10. The transcriptional responses of inoculated synthetic communities to pathogen.** Volcano plots illustrated the differentially expressed genes (DEGs) between pathogen-infected (FC group) and non-infected (C group) plants. RNA and DNA were co-extracted from the same plant sample using a column-based extraction kit. Differential analyses were assessed using DESeq2 with log-transformed DNA counts from the same plant used as an offset for normalization. Dashed lines indicated significance thresholds with $|\log_2 FC| > 2$ and p-value $\leqslant 0.01$. Colored dots represent significantly DEGs with colors corresponding to their reference genomes. The differential analyses were evaluated across three SynCom treatment on day 10. **(a)** SynCom10 **(b)** SynCom11 **(c)** SynCom11_P3.

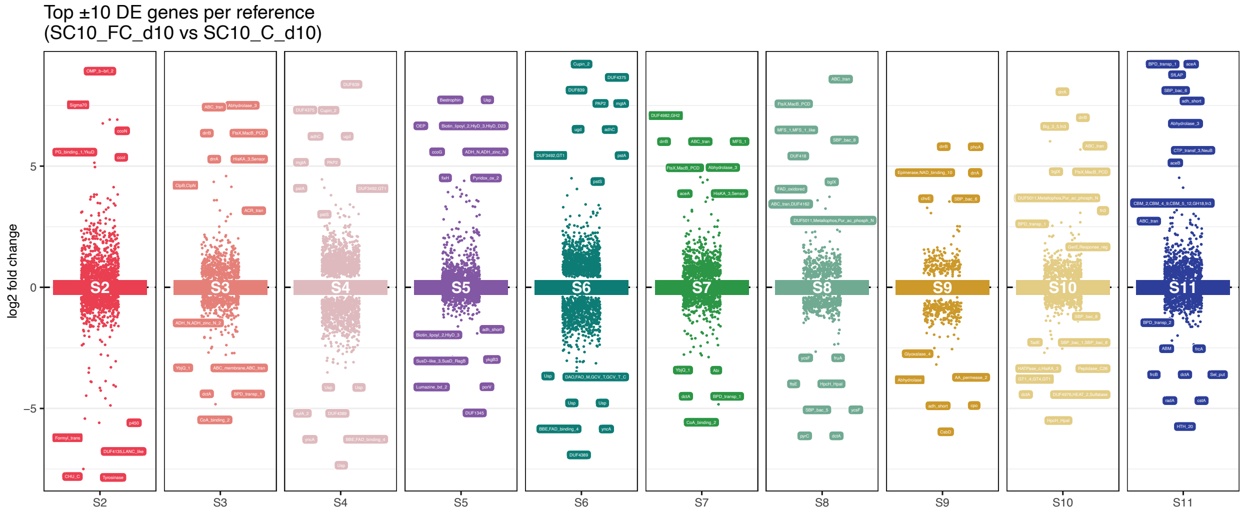


**Fig.S11. The detailed DEGs information of SynCom10 group on day 10.** Different colors corresponding to each reference genomes. The text tags marked the top 10 DEGs per reference that induced by pathogen stress. The y-axis shows the log2 fold change value of these DEGs.

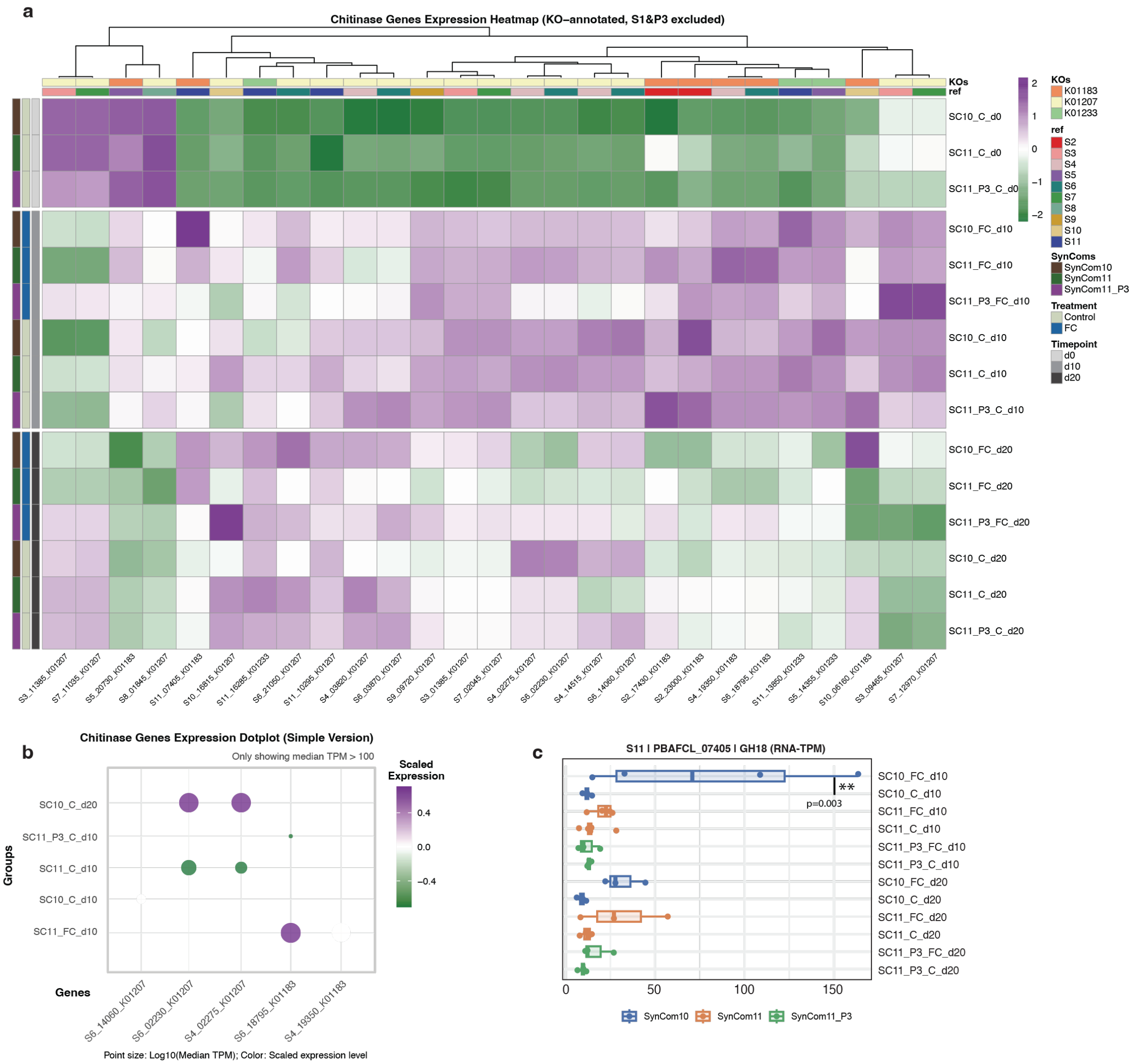


**Fig.S12. The expression overview of chitinase related genes. (a)** The heatmap of all genes that related to chitinase activities based on their KO annotations. The color bar indicated their treatments, reference genome, and KO annotations. The color scale was scaled by per gene within different samples to show the differences of the expression. **(b)** The heatmap dotplot shows the expression of highly activated genes (RNA TPM > 100 in at least one group) that associated with chitinase. The y-axis indicated their treatment groups, and x-axis are the gene information including which genome and their gene position in that genome. **(c)** The expression level of pathogen-induced chitinase gene in the strain S11 (gene: PBAFCL_07405) across all SynComs treatments.

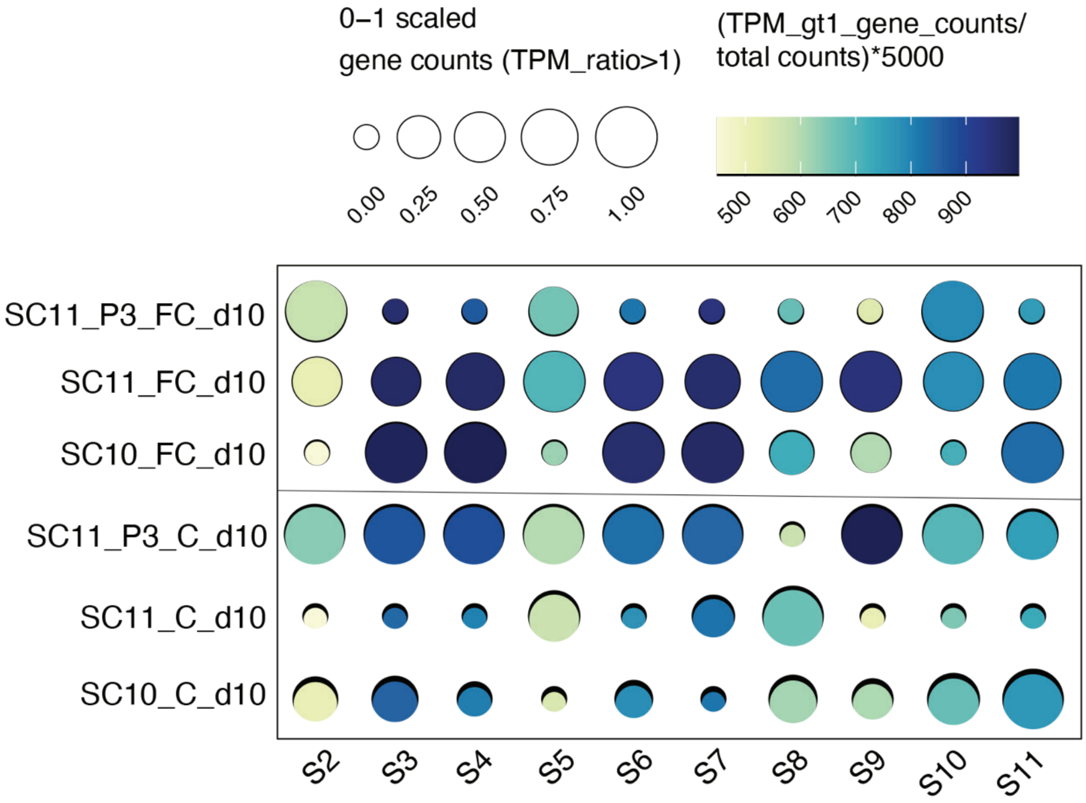


**Fig.S13. The overall gene expression of day 10 samples with/without pathogen inoculation.** The dot size represents the scaled gene counts with a TPM ratio greater than 1 (RNA TPM > DNA TPM). The dot color represents how many genes were activated (RNA TPM > DNA TPM) per 5000 genes. Resource data see Table. S8.

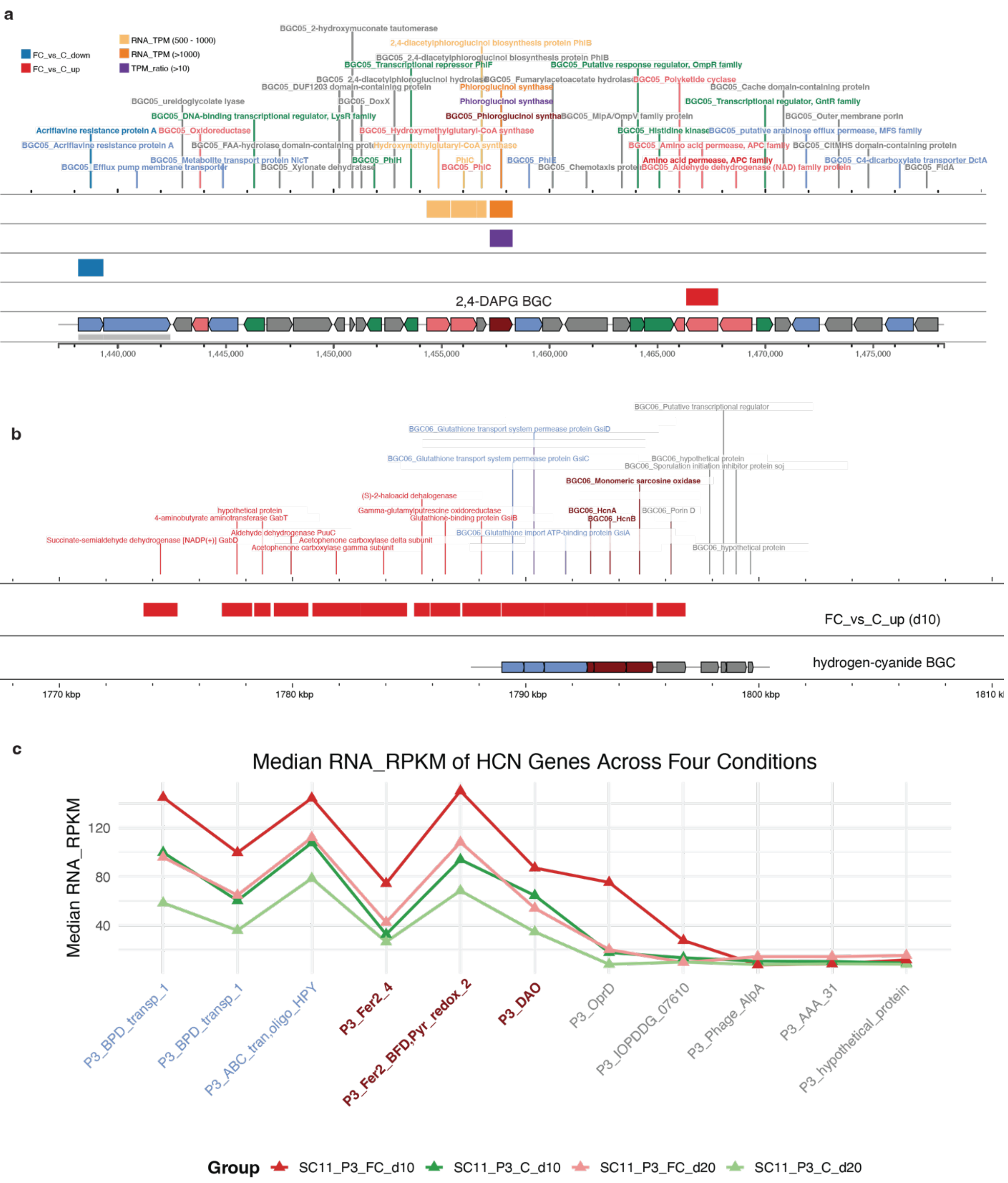


**Fig.S14. The expression overview of the 2,4-DAPG BGC and hydrogen-cyanide (HCN) BGC in strain P3 in SynCom11_P3 treated plants. (a)** The structure of 2,4-DAPG biosynthetic gene cluster and corresponding gene expression pattern. The blue color indicated genes down-regulated under pathogen stress, and red color shows the up-regulated DEGs. The orange color highlighted the DEGs with high absolute transcriptional activity (with RNA TPM greater than 500). The purple color indicates the core genes of this BGC has a TPM ratio greater than 10. **(b)** The structure of HCN BGCs and the red colors indicated the DEGs that upregulated with pathogen pressure. **(c)** The expression of HCN BGC across four conditions of SynCom11_P3 treatment. The expression level is based on median value of RNA RPKM in the samples of the same group. The color indicated which groups it belongs to.

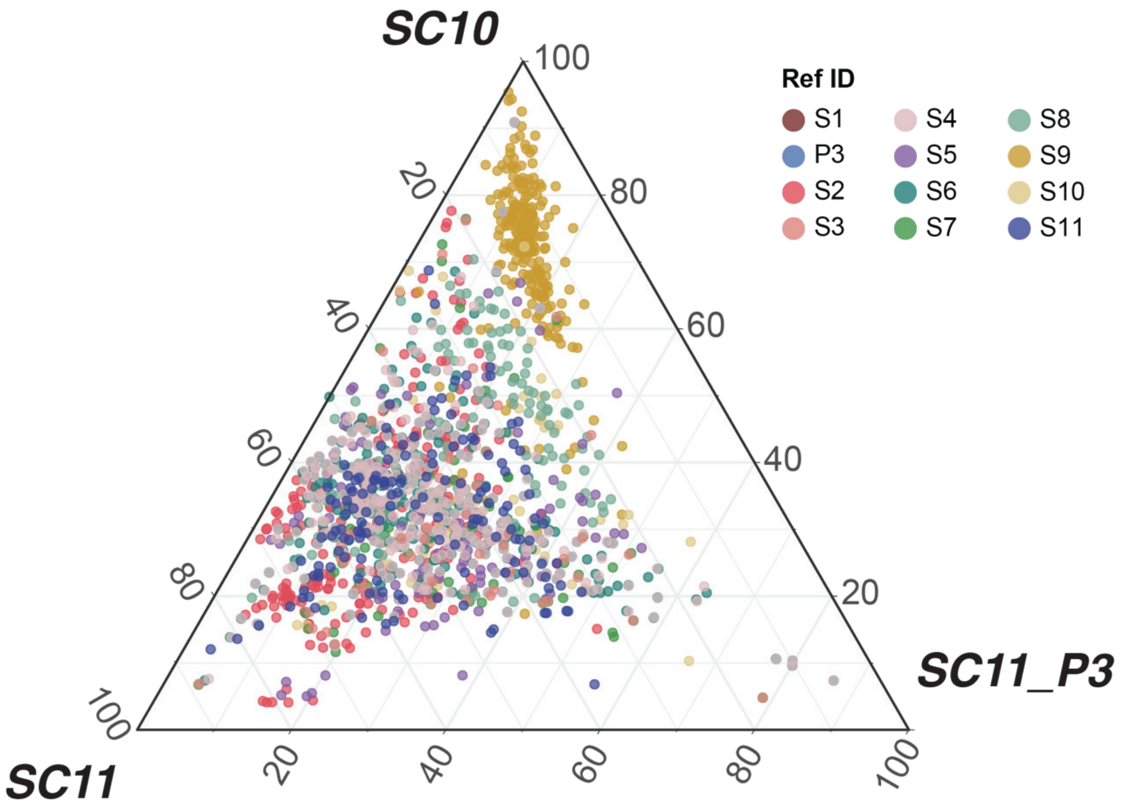


**Fig.S15. Ternary plots of selected DEGs.** These DEGs were up-regulated in at least one SynCom group (FC group versus C group, p-adj ≤ 0.01, $log_2FC > 2$) and activated at day 10 compared to day 20 (within FC group, p-adj ≤ 0.01, $log_2FC > 2$). Gene colors correspond to the strain assignment. Genes from strain S1 and P3 were excluded since they were only present in one of the SynCom.

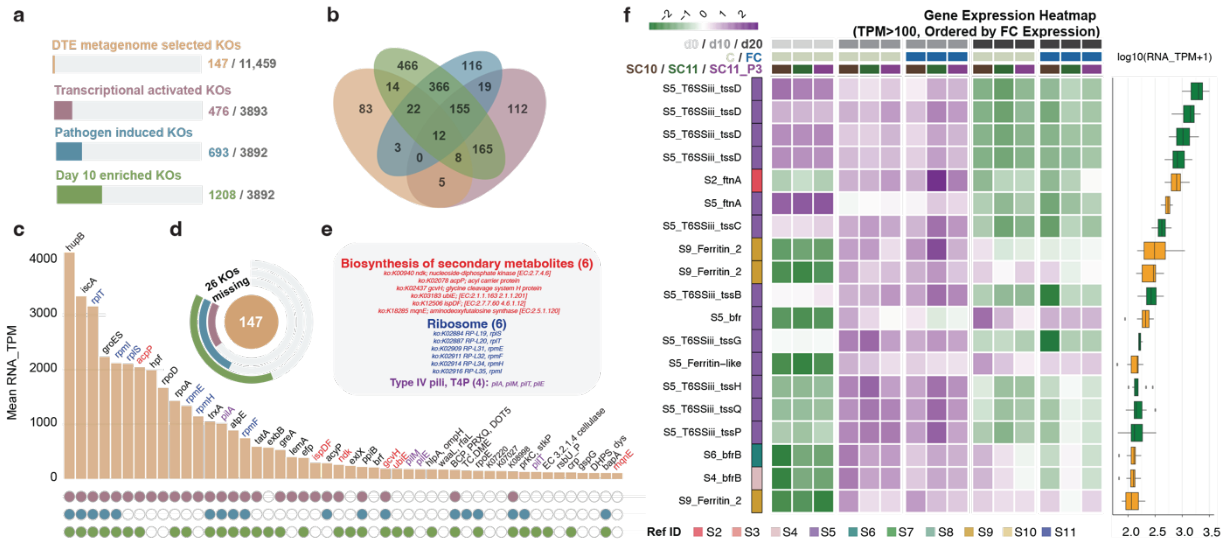

**Fig. S16. The gene expression of suppressive-related functions and pathways in the suppressive SynCom rhizosphere. (a-e)** The combined plots demonstrate the reproducibility of prioritized KOs across disease-suppressive synthetic communities from multiple dimensions. Orange color indicates all KOs statistically enriched in suppressive samples compare to non-suppressive ones in DTE assay. Purple color shows KOs that were highly transcriptionally active (median RNA TPM > 100) in the pathogen- and SynCom-treated rhizosphere microbiome at day 10. Blue color indicates the KOs that were differentially expressed under pathogen stress in SynCom treatments. Green color represents the KOs that were higher expressed in day 10 compared to day 0 and day 20. **Specifically:** a) The overall portions of different categories. b) Venn plot shows the overlapped and unique KOs in these 4 categories. c) Barplot of the top 50 KOs with highest expression levels among 147 suppressive-associated KOs. Colored dots below bars indicate assignment to specific categories d) The donut showing the overlap between DTE-selected KOs and the other three categories. e) Functional annotations of highlighted KOs. **(f)** The heatmap shows the expression level of genes related to iron-uptake (siderophore BGCs, ferritin, ferritin-like, and TonB genes), T6SS, chitinase in different groups. The bottom color bar shows the sample information including date, pathogen stress, and SynCom treatment. The boxplot shows the expression range of each gene and their functional assignment. This plot is based on *Fusarium*-inoculated group at day 10.

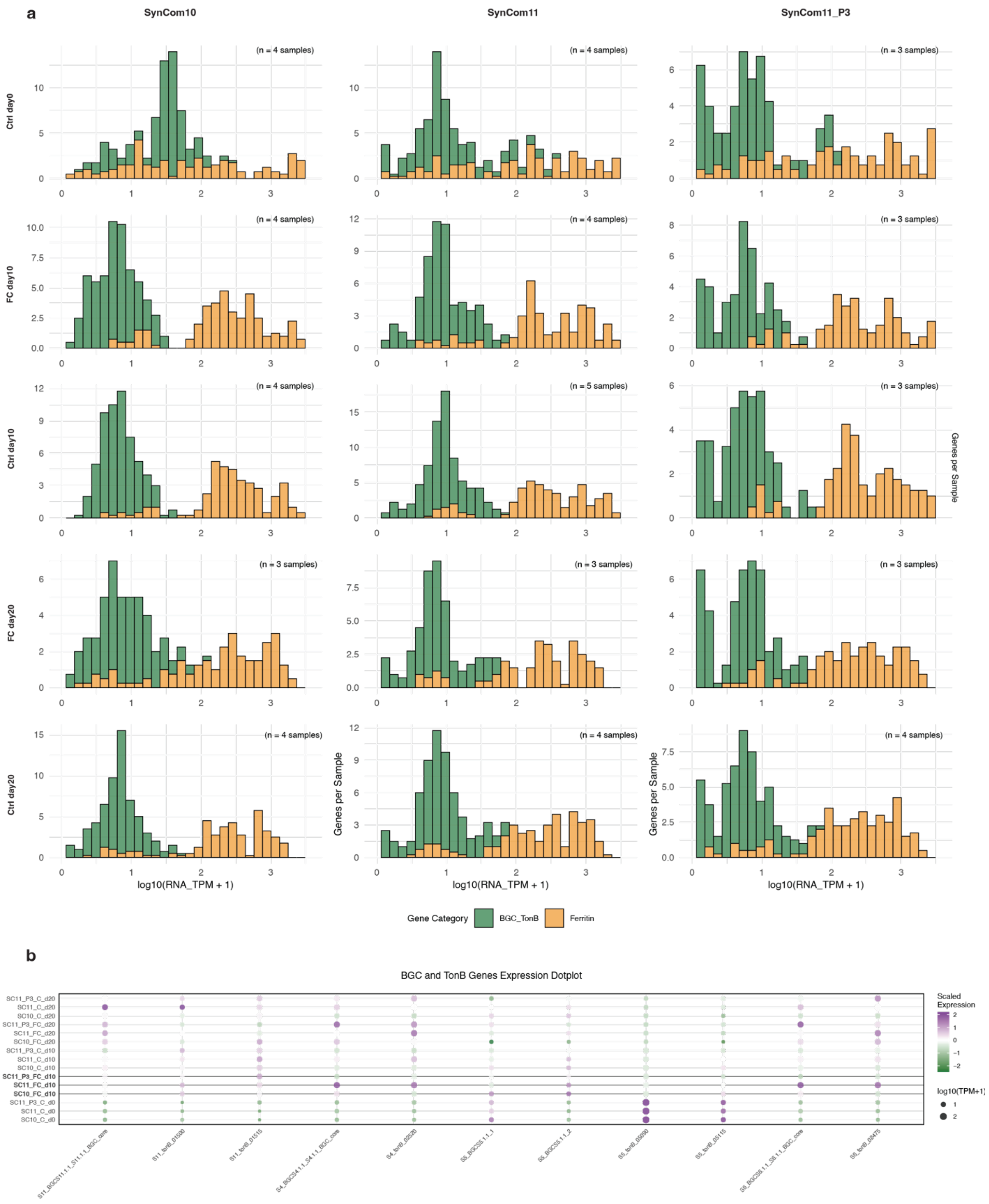


**Fig.S17. The overview of siderophore related genes expression. (a)** The histograms show the expression distribution of siderophore BGC core genes with their TonB genes close to each BGC (gene window size = 50), and the ferrintin-related genes according to their eggnog annotation. There are 56 siderophore-BGC-associated genes and 39 ferrintin-associated genes in the inoculated bacteria genomes. **(b)** The dotplot heatmap shows the expression of highly-activated (with RNA TPM > 100) BGC core genes and TonB genes. The bold font and black line highlighted the day 10 pathogen-infected groups.

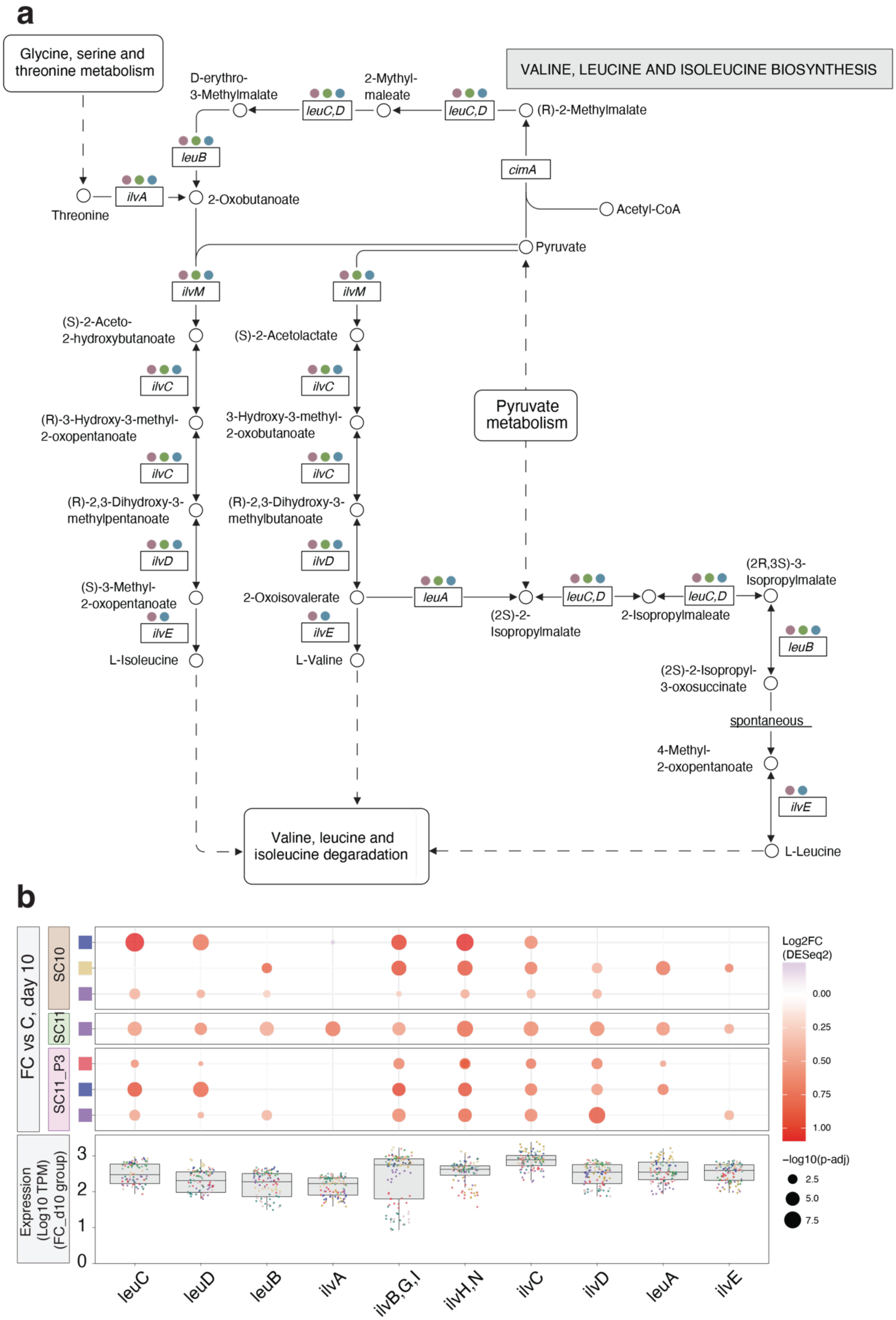
a
Glycine, serine and threonine metabolism
VALINE, LEUCINE AND ISOLEUCINE BIOSYNTHESIS
D-erythro-3-Methylmalate
2-Mythyl-maleate
(R)-2-Methylmalate
leuC,D
leuB
cimA
Acetyl-CoA
Threonine
ilvA
2-Oxobutanoate
Pyruvate
ilvM
(S)-2-Aceto-2-hydroxybutanoate
(S)-2-Acetolactate
ilvC
(R)-3-Hydroxy-3-methyl-2-oxopentanoate
3-Hydroxy-3-methyl-2-oxobutanoate
Pyruvate metabolism
(R)-2,3-Dihydroxy-3-methylpentanoate
(R)-2,3-Dihydroxy-3-methylbutanoate
ilvD
(S)-3-Methyl-2-oxopentanoate
2-Oxoisovalerate
leuA
(2S)-2-Isopropylmalate
2-Isopropylmaleate
(2R,3S)-3-Isopropylmalate
ilvE
L-Isoleucine
L-Valine
(2S)-2-Isopropyl-3-oxosuccinate
spontaneous
4-Methyl-2-oxopentanoate
Valine, leucine and isoleucine degaradation
L-Leucine
b
FC vs C, day 10
SC10
SC11
SC11_P3
Expression (Log10 TPM) (FC_d10 group)
0
1
2
3
leuC
leuD
leuB
ilvA
ilvB,G,I
ilvH,N
ilvC
ilvD
leuA
ilvE
Log2FC (DESeq2)
0.00
0.25
0.50
0.75
1.00
−log10(p-adj)
2.5
5.0
7.5

**Fig.S18. The expression overview of branch-chain amino acid (BCAA) biosynthesis pathway. (a)** The illustrated BCAA biosynthesis pathway (valine, leucine and isoleucine biosynthesis, M00019). The colored dots on top of each KOs in this pathway shows how these KOs are assigned to different layers. **(b)** The heatmap dotplot indicated the pathogen induced genes in BCAA-biosynthesis pathway (M00019). Only the significant genes were shown in the plot. The focused bacteria are the ones that contains more than 5 DEGs in this pathway. The boxplot shows the expression level (displayed in Log10-scale) of genes involved in M00019 pathway in *Fusarium*-inoculated group at day 10. The dot colored by their reference genomes.

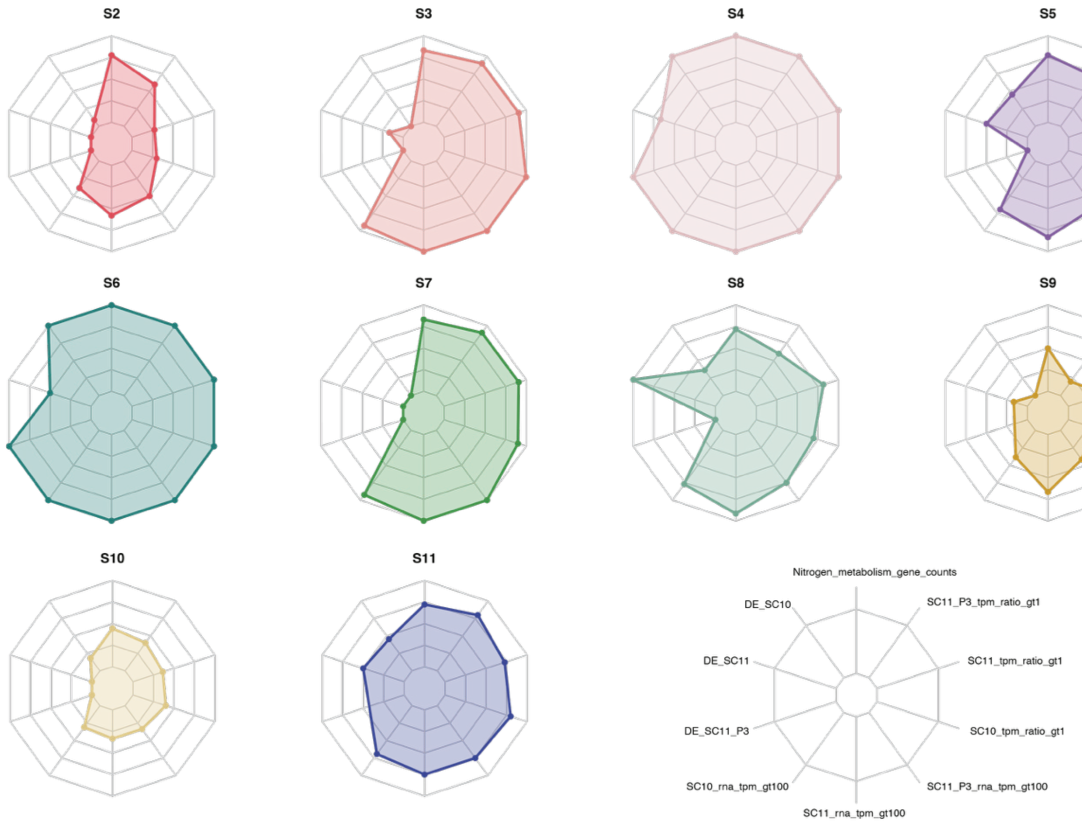


**Fig.S19. The genes on nitrogen metabolisms pathway overview.** Radar plots showing multi-dimensional profiles for different bacterial genomes. Each polygon represents the relative values across 10 parameters for individual genomes. The abbreviations: DE = differentially expressed genes (FC group versus C group, day 10); SC10 = SynCom10; SC11 = SynCom11; SC11_P3 = SynCom11_P3; rna_tpm_gt100 = RNA TPM greater than 100; tpm_ratio_gt1 = RNA TPM / DNA TPM greater than 1.

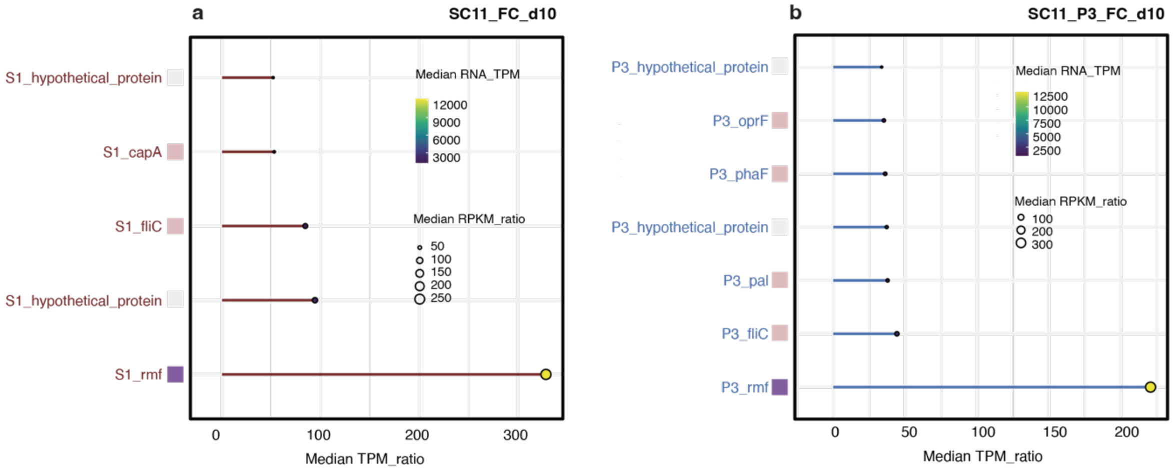


**Fig.S20. The highly expressed genes involved in S1 and P3 genomes.** The lollipop plot is the same illustration setting as Fig.5d. The 2 panels show the results from **(a)** S1 genomes in SynCom11-inoculated pathogen-infected group **(b)** P3 genomes in SynCom11_P3-inoculated pathogen-infected group.

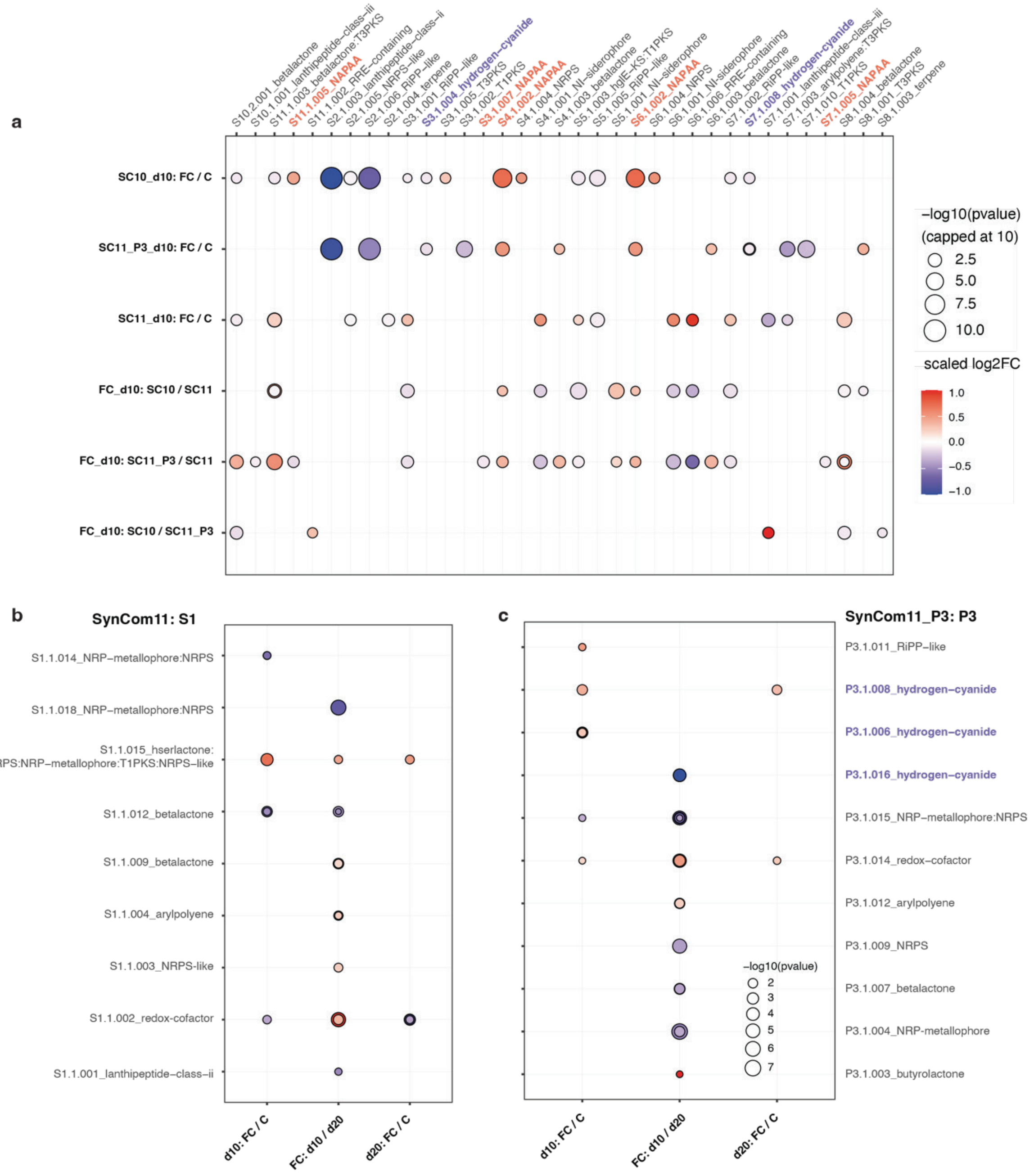


**Fig.S21. The complete overview of expressed BGCs in 3 SynComs.** The heatmap dotplot shows the differentially expression genes (p-adj < 0.01, Log2FC >1, global expression) that belong to BGC core genes. The differential analyses were done in different comparison pairs. BGC that encode non-alpha-poly-amino-acid (NAPAA) and hydrogen-cyanide (HCN) were highlighted in pink and purple font colors. **(a)** BGC from S2 – S11 genomes in different comparison pairs. Y-axis indicate the comparison is between which two groups, the dot size indicate the significance of the differences, and the dot color reflect the log2FC of each core genes. **(b)** BGCs from the S1 strain for SynCom_11 groups. The x-axis shows the comparison groups, and the y-axis shows the BGC type annotations. **(c)** BGCs from the P3 strain for SynCom11_P3 groups. The x-axis shows the comparison groups, and the y-axis shows the BGC type annotations.

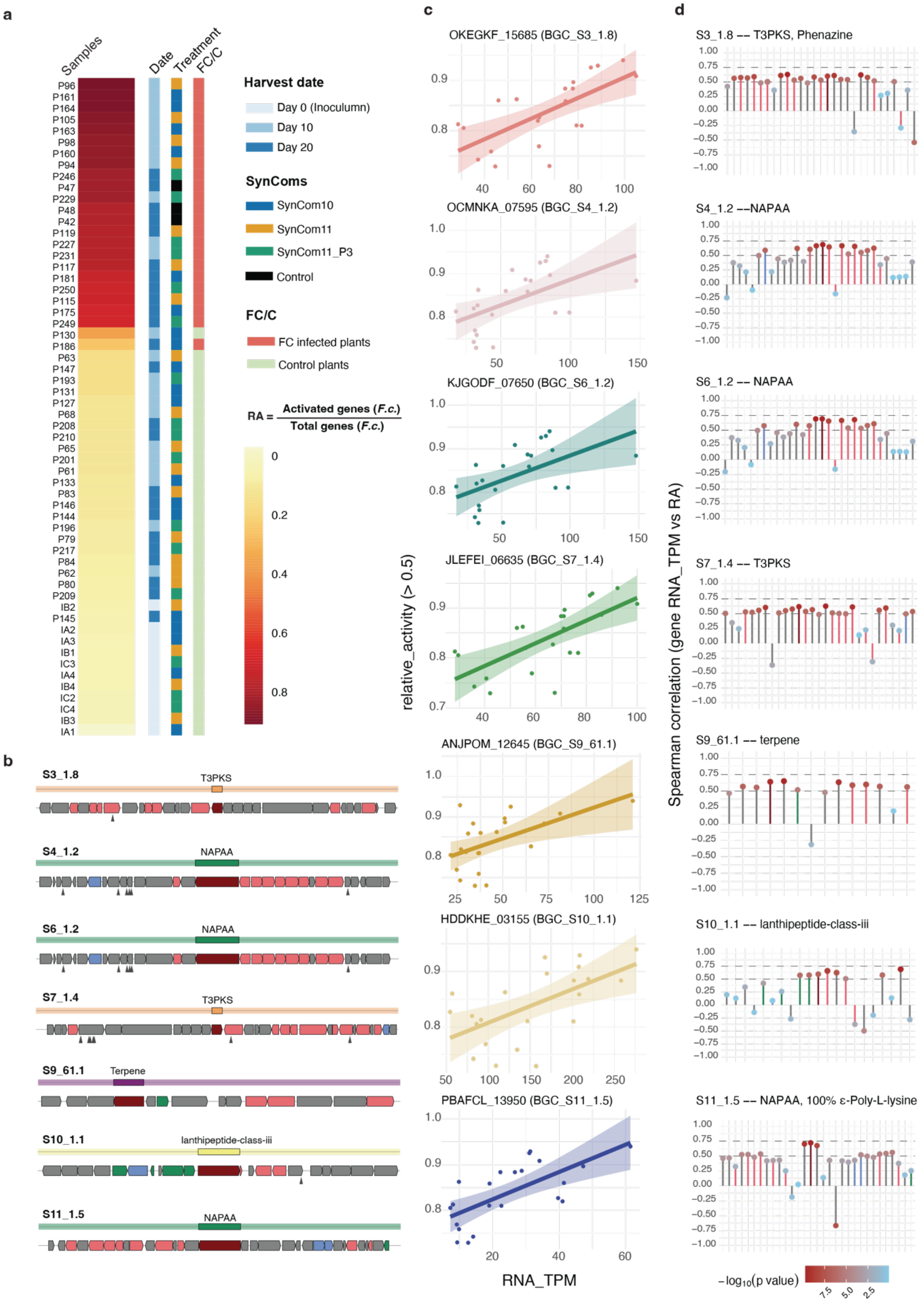
a
Samples
Date
Treatment
FC/C
P96
P161
P164
P105
P163
P98
P160
P94
P246
P47
P229
P48
P42
P119
P227
P231
P117
P181
P250
P115
P175
P249
P130
P186
P63
P147
P193
P131
P127
P68
P208
P210
P65
P201
P61
P133
P83
P146
P144
P196
P79
P217
P84
P62
P80
P209
IB2
P145
IA2
IA3
IB1
IC3
IA4
IB4
IC2
IC4
IB3
IA1
Harvest date
Day 0 (Inoculumn)
Day 10
Day 20
SynComs
SynCom10
SynCom11
SynCom11_P3
Control
FC/C
FC infected plants
Control plants
RA = Activated genes (F.c.) / Total genes (F.c.)
0
0.2
0.4
0.6
0.8
b
S3_1.8
T3PKS
S4_1.2
NAPAA
S6_1.2
NAPAA
S7_1.4
T3PKS
S9_61.1
Terpene
S10_1.1
lanthipeptide-class-iii
S11_1.5
NAPAA
c
OKEGKF_15685 (BGC_S3_1.8)
OCMNKA_07595 (BGC_S4_1.2)
KJGODF_07650 (BGC_S6_1.2)
JLEFEI_06635 (BGC_S7_1.4)
ANJPOM_12645 (BGC_S9_61.1)
HDDKHE_03155 (BGC_S10_1.1)
PBAFCL_13950 (BGC_S11_1.5)
relative_activity (> 0.5)
RNA_TPM
d
S3_1.8 -- T3PKS, Phenazine
S4_1.2 --NAPAA
S6_1.2 -- NAPAA
S7_1.4 -- T3PKS
S9_61.1 -- terpene
S10_1.1 -- lanthipeptide-class-iii
S11_1.5 -- NAPAA, 100% ε-Poly-L-lysine
Spearman correlation (gene RNA_TPM vs RA)
−log10(p value)
7.5
5.0
2.5

**Fig.S22. The overview of biosynthetic gene clusters that shows linear positive correlation with pathogen activities. (a)** The color scale represents relative activities of *F. culmorum* in each sample. The color bar next to it shows their group information including SynComs-treatment, pathogen stress, and their harvest date. We use the activated genes (TPM ratio >10) numbers divided by total genes of *F. culmorum* genome to define a relative activity for the correlation analysis. **(b)** The selected BGCs that have core genes with a linear positive correlation with pathogen activities in pathogen-inoculated samples. **(c)** The linear model plots show the correlation between the core gene expression and the pathogen relative activities. **(d)** The correlation of all genes involved in each BGC. The y-axis is the correlation spearmanr-value. The dot color indicates the -log10(p-adj) value that shows the significance of the correlations.

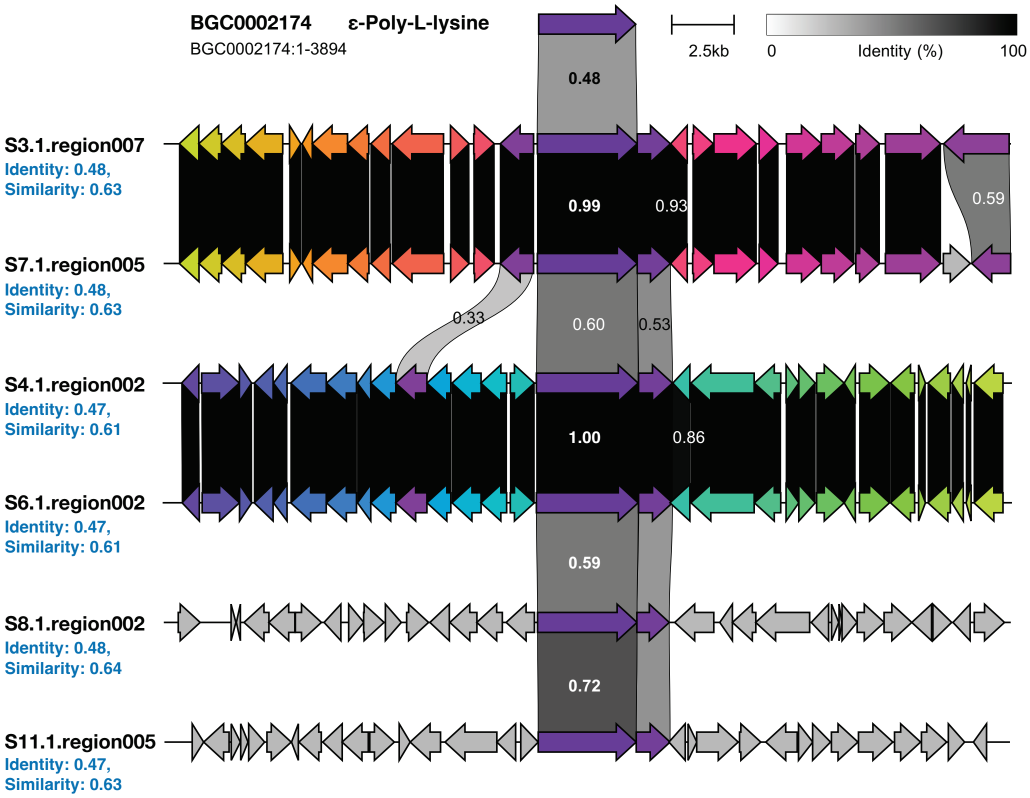


**Fig.S23. The overview of NAPAA biosynthetic gene clusters in bacterial candidates for SynCom assay.** The number on the color bar indicate the identity of the gene pairs. The identity value of gene pairs with more than 0.98 identity is hide. The identity and similarity of core gene compare to epsilon-poly-L-lysine gene is marked with blue color.

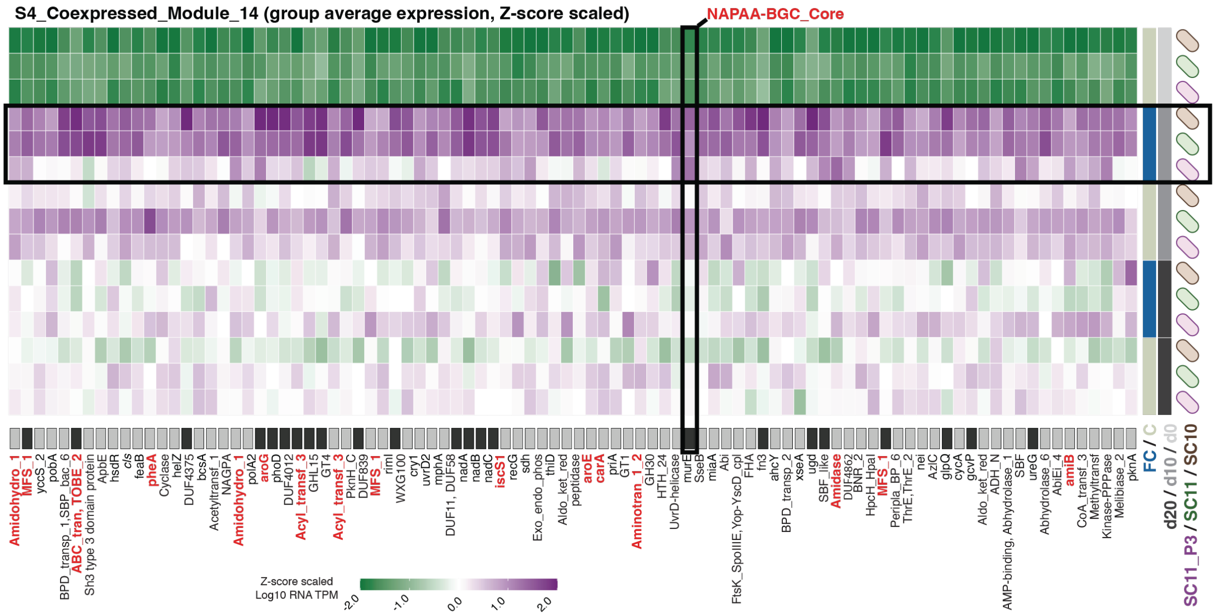


**Fig.S24. The heatmap shows the expression level of S4_Coexpressed_Module_14 in different groups.** The core gene of S4_NAPAA BGC were highlighted in a black rectangle. The genes related to amino acid biosynthesis, metabolism, and transport were highlighted in red color. The color bar on the right side shows the different conditions of each group with pathogen stress, SynComs, and harvest date. The black boxes under the heatmap indicated the DEGs that up-regulated with pathogen stress in at least one SynCom treatment at day 10.

#### References


1. Tracanna, V. *et al.* Dissecting Disease-Suppressive Rhizosphere Microbiomes by Functional Amplicon Sequencing and 10× Metagenomics. *mSystems* **6**, e01116-20 (2021).
2. Lemanceau, P., Expert, D., Gaymard, F., Bakker, P. A. H. M. & Briat, J.-F. Chapter 12 Role of Iron in Plant–Microbe Interactions. in *Advances in Botanical Research* vol. 51 491–549 (Elsevier, 2009).
3. Mazurier, S., Corberand, T., Lemanceau, P. & Raaijmakers, J. M. Phenazine antibiotics produced by fluorescent pseudomonads contribute to natural soil suppressiveness to Fusarium wilt. *The ISME Journal* **3**, 977–991 (2009).

4. Siegel-Hertz, K. *et al.* Comparative Microbiome Analysis of a Fusarium Wilt Suppressive Soil and a Fusarium Wilt Conducive Soil From the Châteaurenard Region. *Front. Microbiol.* **9**, 568 (2018).
5. Carrión, V. J. *et al.* Pathogen-induced activation of disease-suppressive functions in the endophytic root microbiome. *Science* **366**, 606–612 (2019).
6. Coulthurst, S. J. The Type VI secretion system – a widespread and versatile cell targeting system. *Research in Microbiology* **164**, 640–654 (2013).
7. Records, A. R. The Type VI Secretion System: A Multipurpose Delivery System with a Phage-Like Machinery. *MPMI* **24**, 751–757 (2011).
8. Monjarás Feria, J. & Valvano, M. A. An Overview of Anti-Eukaryotic T6SS Effectors. *Front. Cell. Infect. Microbiol.* **10**, 584751 (2020).
9. Trunk, K. *et al.* The type VI secretion system deploys antifungal effectors against microbial competitors. *Nat Microbiol* **3**, 920–931 (2018).
10. Vogel, C. M., Potthoff, D. B., Schäfer, M., Barandun, N. & Vorholt, J. A. Protective role of the Arabidopsis leaf microbiota against a bacterial pathogen. *Nat Microbiol* **6**, 1537–1548 (2021).
11. Chewning, S. S. *et al.* Root-Associated *Streptomyces* Isolates Harboring *melC* Genes Demonstrate Enhanced Plant Colonization. *Phytobiomes Journal* **3**, 165–176 (2019).
12. Alavi, P., Starcher, M. R., Zachow, C., Müller, H. & Berg, G. Root-microbe systems: the effect and mode of interaction of Stress Protecting Agent (SPA) Stenotrophomonas rhizophila DSM14405T. *Front. Plant Sci.* **4**, (2013).

13. McClerklin, S. A. *et al.* Indole-3-acetaldehyde dehydrogenase-dependent auxin synthesis contributes to virulence of Pseudomonas syringae strain DC3000. *PLoS Pathog* **14**, e1006811 (2018).

14. Neuhäuser, B., Dynowski, M., Mayer, M. & Ludewig, U. Regulation of NH4 + Transport by Essential Cross Talk between AMT Monomers through the Carboxyl Tails. *Plant Physiology* **143**, 1651–1659 (2007).

15. Van Der Palen, C. J. N. M., Reijnders, W. N. M., De Vries, S., Duine, J. A. & Van Spanning, R. J. M. MauE and MauD proteins are essential in methylamine metabolism of Paracoccus denitrificans. *Antonie Van Leeuwenhoek* **72**, 219–228 (1997).

16. Jurgenson, C. T., Begley, T. P. & Ealick, S. E. The Structural and Biochemical Foundations of Thiamin Biosynthesis. *Annu. Rev. Biochem.* **78**, 569–603 (2009).

17. Izumi, A., Schnell, R. & Schneider, G. Crystal structure of NirD, the small subunit of the nitrite reductase NirbD from *Mycobacterium tuberculosis* at 2.0 Å resolution. *Proteins* **80**, 2799–2803 (2012).

18. Kuzyakov, Y. & Xu, X. Competition between roots and microorganisms for nitrogen: mechanisms and ecological relevance. *New Phytologist* **198**, 656–669 (2013).

19. Pandit, M. A. *et al.* Major Biological Control Strategies for Plant Pathogens. *Pathogens* **11**, 273 (2022).

20. Tripathi, R. *et al.* Plant mineral nutrition and disease resistance: A significant linkage for sustainable crop protection. *Front. Plant Sci.* **13**, 883970 (2022).

21. Singh, S. S. *et al.* Brevicillin, a novel lanthipeptide from the genus *Brevibacillus* with antimicrobial, antifungal, and antiviral activity. *Journal of Applied Microbiology* **134**, lxad054 (2023).

22. Clough, S. E., Jousset, A., Elphinstone, J. G. & Friman, V. Combining in vitro and in vivo screening to identify efficient *Pseudomonas* biocontrol strains against the phytopathogenic bacterium *Ralstonia solanacearum*. *MicrobiologyOpen* **11**, e1283 (2022).
23. Dutta, S., Yu, S.-M. & Lee, Y. H. Assessment of the Contribution of Antagonistic Secondary Metabolites to the Antifungal and Biocontrol Activities of Pseudomonas fluorescens NBC275. *Plant Pathol J* **36**, 491–496 (2020).
24. Kim, D.-R. & Kwak, Y.-S. A Genome-Wide Analysis of Antibiotic Producing Genes in Streptomyces globisporus SP6C4. *Plant Pathol J* **37**, 389–395 (2021).
25. Kim, D.-R., Jeon, C.-W. & Kwak, Y.-S. Antifungal Properties of Streptomyces bacillaris S8 for Biological Control Applications. *Plant Pathol J* **40**, 322–328 (2024).
26. Chu, D. *et al.* Genomic insights on fighting bacterial wilt by a novel *Bacillus amyloliquefaciens* strain Cas02. *Microbial Biotechnology* **15**, 1152–1167 (2022).
27. Yang, F. *et al.* Mechanism of a novel Bacillus subtilis JNF2 in suppressing Fusarium oxysporum f. sp. cucumerium and enhancing cucumber growth. *Front. Microbiol.* **15**, 1459906 (2024).
28. Heo, Y., Lee, Y., Balaraju, K. & Jeon, Y. Characterization and evaluation of Bacillus subtilis GYUN-2311 as a biocontrol agent against Colletotrichum spp. on apple and hot pepper in Korea. *Front. Microbiol.* **14**, 1322641 (2024).
29. Abdalla Abdelshafy Mohamad, O. *et al.* Dual-functionality of Nocardiopsis alba B57 in biocontrol and plant growth: a metabolomic approach to agricultural sustainability. *npj Biofilms Microbiomes* **11**, 164 (2025).

30. Widada, J., Damayanti, E., Alhakim, M. R., Yuwono, T. & Mustofa, M. Two strains of airborne *Nocardiopsis alba* producing different volatile organic compounds (VOCs) as biofungicide for *Ganoderma boninense*. *FEMS Microbiology Letters* **368**, fnab138 (2021).
